# YhdP forms an envelope-spanning phospholipid-binding bridge in complex with DedD

**DOI:** 10.64898/2026.10.30.763061

**Authors:** Qiaoyu Tian, Robert Clark, Isabella Musto, Carla Kirschbaum, Jamie Mitra, Benjamin F. Cooper, Joanna Szczepaniak, Marcie Bamber, Carol V. Robinson, Syma Khalid, Georgia L. Isom

## Abstract

The outer membrane of Gram-negative bacteria is built from phospholipids synthesised at the inner membrane, but how they cross the periplasm remains unresolved. YhdP is the leading candidate for anterograde phospholipid transfer and is predicted to form an envelope-spanning bridge. Here we purify YhdP at endogenous levels and show by cryo-electron microscopy that it contains a continuous cavity, alongside an additional density that proteomics identifies as the cell wall-binding protein DedD. Liposome co-sedimentation shows that the C-terminal helices of YhdP bind membranes directly, and single-molecule tracking reveals that endogenous YhdP is as immobile as an outer membrane protein, indicating an envelope-spanning architecture *in vivo*. Native mass spectrometry demonstrates that the YhdP groove accommodates all major classes of *E. coli* phospholipid without headgroup preference. DedD engages YhdP through its disordered region and increases lipid density at the groove entrance, suggesting a potential role in lipid loading.

## Introduction

Antibiotic-resistant Gram-negative bacteria pose a severe threat to global healthcare^1^. Gram-negative bacteria differ from Gram-positive bacteria mainly due to a thinner peptidoglycan layer and the presence of an outer membrane (OM). The OM consists of an outer leaflet of lipopolysaccharides (LPS) and an inner leaflet of phospholipids, with embedded OM proteins (OMPs) and lipoproteins. Maintenance of the lipid asymmetry in the OM is important for bacteria to defend against antimicrobials^2,3^. All components of the OM are synthesised and modified in the cytoplasm and the inner membrane (IM), meaning that they require transport across the hydrophilic periplasm to the OM^3^. For LPS, OMPs, and lipoproteins, the transportation machinery has been well characterised^4–6^. In contrast, the mechanisms of how phospholipids get transported from the IM to OM remains poorly understood.

More recently, the AsmA-like protein family has been implicated in this anterograde phospholipid transfer^7–9^. There are six family members in *E. coli*, namely YhdP, TamB, YdbH, AsmA, YicH and YhjG, among which YhdP is the most strongly implicated in phospholipid transfer. The link between YhdP and anterograde phospholipid transfer was first discovered in a genetic screen of *E. coli* harbouring the dominant negative *mlaA*\* mutation^7^. The destablised OM of the *mlaA\** strain results in excessive shedding of OM material, followed by subsequent shrinkage of the IM as lipids are transferred to replenish the OM^7^. Deletion of *yhdP* in this *mlaA*\* strain slows down the IM shrinkage, indicating a reduction of anterograde phospholipid transfer. In support of a role in OM homeostasis, a *yhdP* knockout strain is sensitive to polymyxin, SDS+EDTA and vancomycin^7,8,10–12^. Furthermore, deletion of *tamB* and *ydbH* in a *yhdP* knockout strain results in loss of cell viability, revealing that these three largest AsmA-like family members are collectively essential^8,11^. Decreasing expression of *yhdP* in a Δ*tamB*Δ*ydbH* background leads to an accumulation of phospholipids in the IM^8^, suggesting that all three proteins may contribute to transfer of phospholipids to the OM. Recently, purified YhdP and TamB have been shown to aid exchange of lipids between liposomes^13^. Overall, these data suggest that AsmA-like protein family members, in particular YhdP, play an important role in phospholipid transfer and OM homeostasis.

AlphaFold models have provided molecular insights into how the AsmA-like proteins may function. All six AsmA-like family proteins have a single-pass transmembrane helix at the IM and a periplasmic domain made of stacked β sheets of varying lengths^9^. The periplasmic domain of each family member is predicted to have a hydrophobic β-groove along its length that could shield the hydrophobic acyl chains of phospholipids from the hydrophilic periplasm^9^. Further, our previous *in vivo* crosslinking experiments captured phosphate-containing molecules inside the groove of YhdP^12^, though these are yet to be identified as phospholipids. The three longest members, YhdP, TamB and YdbH, have been shown to span the periplasm and interact with the OM, either directly or via proteins at the OM. TamB and YdbH are known to interact with TamA and YnbE, respectively^14–18^. In contrast, there is no known interactor of YhdP at the OM. In our previous molecular dynamics (MD) simulations, the C-terminal helices of YhdP, which are required for function, are able to directly interact with the OM^12^. Recently, it was reported that the IM-anchored cell wall interacting protein, DedD, binds YhdP^19^. DedD is a SPOR-domain containing protein that localises to denuded peptidoglycan during cell division^20–22^. However, the significance of its interaction with YhdP remains unknown. Together, these data support a model that YhdP directly spans from the IM to the OM to transfer phospholipids. However, many aspects of this model are yet to be validated. There are no experimental structures of YhdP, and the role of DedD in YhdP function remains unknown. Further, there is no experimental data to support the C-terminus of YhdP interacting with membranes, nor have the phosphate-containing molecules captured in the groove been identified as phospholipids.

Here, we obtain low resolution cryo-EM data of endogenously purified YhdP, and confirm the presence of a continuous cavity along its length. Liposome co-sedimentation assays support the model that YhdP forms an envelope-spanning groove that directly interacts with the OM, and native mass spectrometry reveals YhdP can accommodate all three main classes of *E. coli* phospholipid. In agreement with a recent study^19^, DedD is identified as an interactor of YhdP. The molecular basis of this interaction is probed by mutagenesis, revealing the disordered domain of DedD is key for the interaction. The localisation of YhdP and DedD at endogenous levels are studied by super-resolution light microscopy, revealing DedD does not obviously recruit YhdP to the septum during cell division. Instead, MD simulations suggest that the role of DedD is to increase lipid density at the N-terminus of YhdP, potentially to enhance lipid loading into the YhdP groove for transfer to the OM.

## Results

### Endogenous YhdP contains a continuous cavity and binds DedD

To capture YhdP in its most native state to study its structure and interactors, we aimed to purify endogenous levels of YhdP from *E. coli* for single-particle cryogenic electron microscopy (cryo-EM) and proteomics. A Twin-Strep-tag was inserted endogenously at the N- or C-terminus of *yhdP* in *E. coli* K-12 BW25113 via lambda red recombination^23,24^. To test if chromosomal tagging of YhdP had impaired its function, growth of both N and C-terminally tagged strains were compared to *ΔyhdP*, which has known sensitivity to SDS+EDTA^7,8,11,12^. The N-terminally tagged strain grew similarly to the *ΔyhdP* strain, suggesting that the N-terminal tag did impair YhdP function (Fig. S1A,B). In contrast, the C-terminally tagged strain grew like WT *E. coli* under these conditions, suggesting YhdP remained functional (Fig. S1A,B).

Endogenous C-terminally tagged YhdP was purified (Fig. S1C) and plunge frozen on 2nm carbon coated copper grids for single particle cryo-EM data collection and processing (Fig. S2). 2D classes revealed that YhdP aligned the best at its periplasmic domain, with the N-terminus and C-terminus less clear, probably due to flexibility at both ends (Fig. 1A). These 2D classes showed a continuous clear cavity throughout the middle, which curved with subtle kinks at several locations.

**Figure 1.**
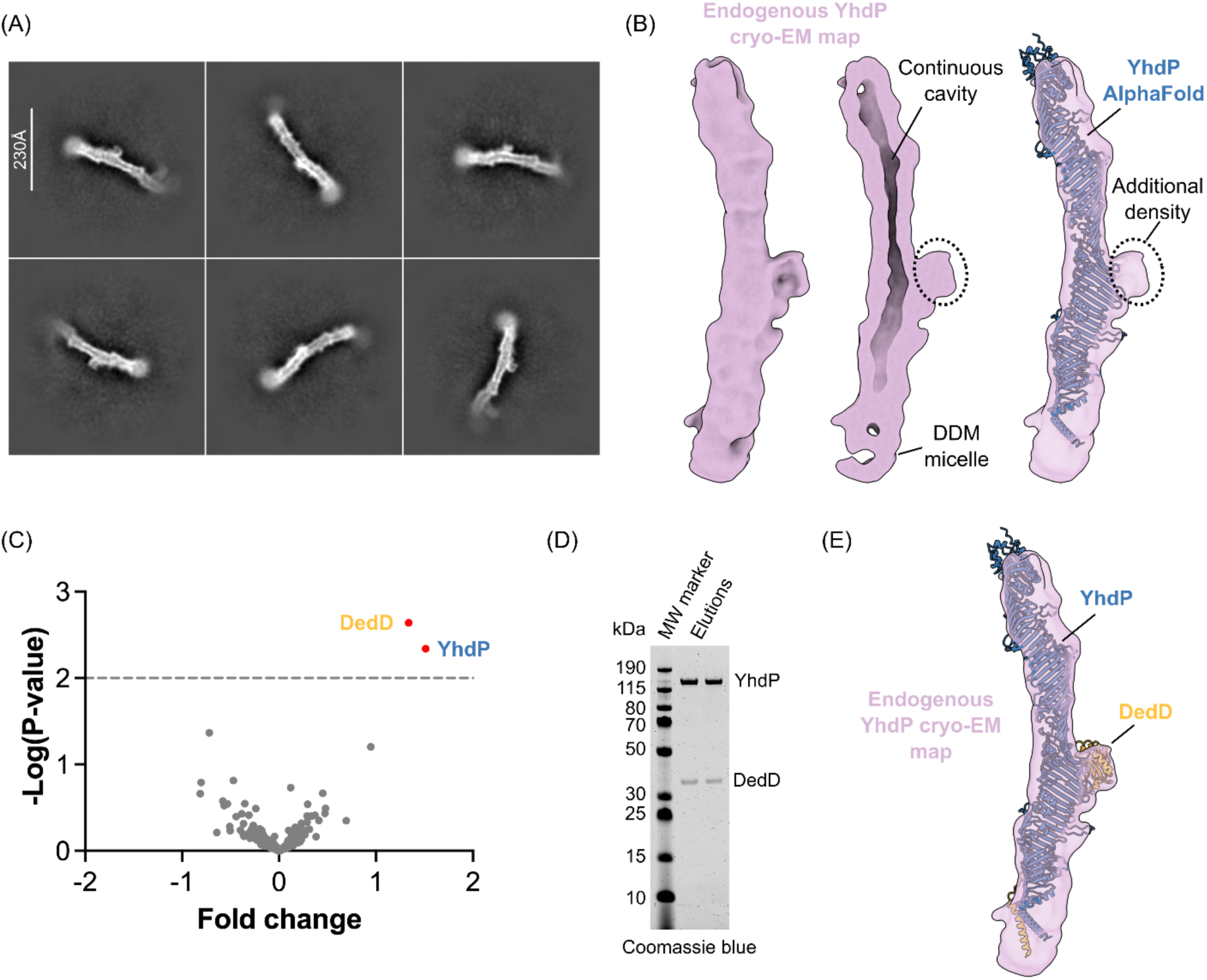
Cryo-EM and proteomics of endogenous YhdP reveals a continuous cavity and DedD as an interactor. (A) Endogenously purified YhdP was studied by single-particle cryo-EM, with representative 2D class averages shown. (B) Reconstructed 3D volume from cryo-EM coloured in purple (left and middle). The density was clipped through to visualise the continuous cavity (middle). Features, corresponding to additional protein and a potential DDM micelle, are labelled. For comparison, the AlphaFold predicted model of YhdP is shown on the right, coloured in blue, with the location of the additional density labelled. (C) Volcano plot comparing proteins isolated from WT *E. coli* and C-terminally Twin-Strep-tagged YhdP following Strep-affinity purification. Significantly enriched hits are labelled (p<0.1). (D) Representative SDS-PAGE of elution fractions following tandem affinity chromatography purification of the overexpressed YhdP-DedD complex. (E) AlphaFold predicts a YhdP-DedD complex. The YhdP-DedD AlphaFold model is fit into the endogenous YhdP cryo-EM map. pLDDT scores and the PAE plot are shown in Fig. S3A.

A low-resolution 8.3 Å 3D map was reconstructed (Fig. 1B), revealing a continuous cavity along the periplasmic domain of YhdP. Compared to the AlphaFold model of YhdP, an additional density was observed at the middle section of YhdP, potentially revealing an interactor (Fig. 1B). Overall, this EM map provided experimental proof that YhdP has a cavity along its length, and reveals an endogenously purified interactor for further investigation.

To identify the additional density observed in the Cryo-EM map, endogenous C-terminally tagged YhdP was purified alongside an untagged WT control strain, for proteomics. Alongside YhdP, DedD was significantly enriched from the tagged YhdP samples (Fig. 1C), consistent with the recent report of DedD interacting with YhdP^19^. However, the molecular characteristics and role of this interaction remain unknown. DedD is a protein of unknown function that contains an N-terminal transmembrane helix (TMH) and a C-terminal sporulation-related repeat (SPOR) domain connected by an intrinsically disordered region (IDR). The SPOR domain is known to bind to denuded peptidoglycan during cell division^25,20–22,26^. To validate this interaction, YhdP and DedD were tagged with a StrepII tag and a His tag, respectively, and over-expressed together. After tandem affinity chromatography, both proteins could be observed by SDS-PAGE, confirming a YhdP-DedD interaction (Fig. 1D). In agreement, AlphaFold^27^ predicted a model of the YhdP-DedD complex (Fig. 1E, Fig. S3A), where the SPOR domain of DedD docks onto the side of YhdP. The AlphaFold model fitted well into the endogenous YhdP cryo-EM map, indicating that the additional density observed is likely attributed to the SPOR domain of DedD (Fig. 1E). Both the AlphaFold model and our cryo-EM map indicated a 1:1 stoichiometry between YhdP and DedD. This was further confirmed by native MS, which analyses proteins in their native folded state^28,29^ (Fig. S3B). Together, these data identify DedD as a native binding partner of YhdP and indicate that the additional density observed in the endogenous cryo-EM reconstruction likely corresponds to the DedD SPOR domain. These findings establish the YhdP–DedD complex as a physiologically relevant assembly and motivate further biochemical and structural characterisation.

### YhdP spans the cell envelope

To efficiently transfer phospholipids from the IM to the OM, YhdP would need to interact with both membranes. At its N-terminus, YhdP has a transmembrane helix that anchors it to the IM. At the C-terminus, YhdP has two helical loops (C-helix_1: residues 1203-1237, C-helix_2: residues 1121-1144). The YhdP AlphaFold prediction was simulated in a double membrane system using molecular dynamics, where both helical loops embedded in the inner leaflet of the OM (Fig. 2A), consistent with our previous simulations^12^. Furthermore, replacing either of these helical loops with disordered linkers results in non-functional YhdP, confirming that these helical loops are essential^12^.

**Figure 2.**
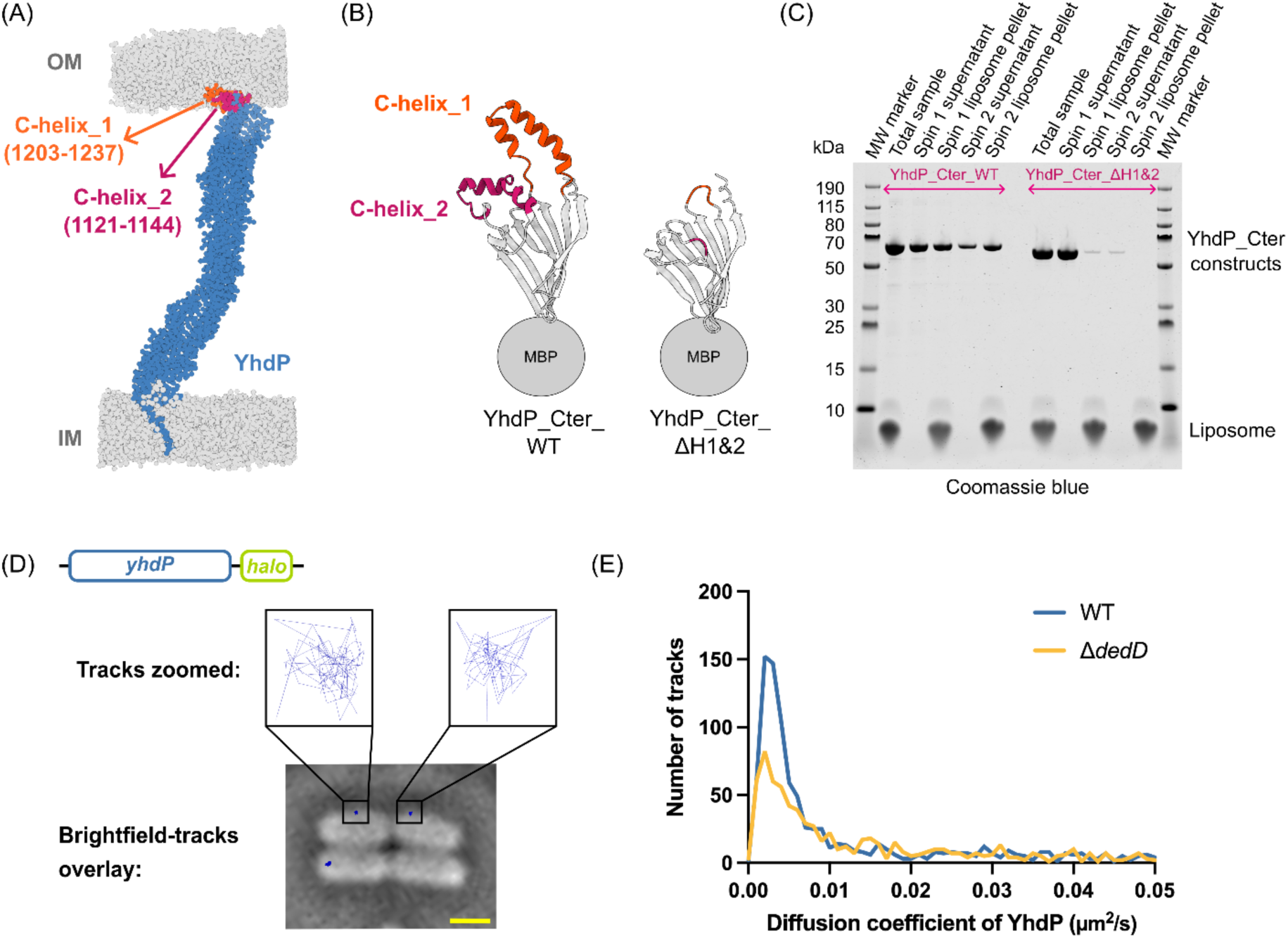
YhdP functions as a cell-envelope spanning bridge. (A) Representative molecular dynamics snapshot of YhdP with both the IM and OM is shown, with the two C-terminal helical loops highlighted as labelled. Beads representing phospholipids or LPS are coloured in grey. During the simulation, both C-terminal helical loops interact with the inner leaflet of the OM. (B,C) Liposome co-sedimentation assay was performed with two C-terminal constructs (YhdP_Cter_WT and YhdP_Cter_ΔH1&2). Schematic diagrams of the two constructs are shown in (B), consisting of AlphaFold predicted models and the N-terminal MBP. Normalised samples across the liposome co-sedimentation assay were analysed by SDS-PAGE, representative images shown in (C). (D) Representative trajectories (blue) from single-molecule tracking of endogenous C-terminally tagged YhdP. A gene schematic of the strain used is shown above. Scale bar (yellow): 1 µm. (E) Histogram of apparent diffusion coefficient calculated from C-terminally Halo tagged YhdP with DedD (WT) and without DedD (Δ*dedD*). Mean±SEM of apparent diffusion coefficient for WT and Δ*dedD* are 0.025±0.0014 µm^2^/s and 0.035±0.0018 µm^2^/s. The apparent diffusion coefficient is calculated with the anomalous diffusion exponent of 1.

To test if these helical loops interact with membranes *in vitro*, membrane association of a C-terminal fragment of YhdP (residues 1101 - 1266, YhdP_Cter_WT, Fig. 2B) was tested by liposome co-sedimentation, an assay that is widely used to study protein-membrane interactions^30,31^. In addition, a negative control where the two C-terminal helical loops were replaced with GS linkers (YhdP_Cter_ΔH1&2, Fig. 2B) was constructed. To aid protein solubility, both fragments were conjugated with MBP at the N-terminus. Both YhdP_Cter_WT and YhdP_Cter_ΔH1&2 proteins were purified as soluble proteins (Fig. S4) and were incubated with liposomes reconstituted from *E. coli* total lipid extract. Liposomes plus any interacting YhdP were pelleted by ultracentrifugation, washed once to remove non-specific binding, and pelleted again. SDS-PAGE revealed that a large proportion of YhdP_Cter_WT co-sedimented with liposomes and that this interaction was maintained during both ultracentrifugation steps (Fig. 2C). In contrast, only a small proportion of YhdP_Cter_ΔH1&2 was observed after the first co-sedimentation, and this was lost after the wash step indicating non-specific binding (Fig. 2C). Collectively, these data reveal that the C-terminal helical loops of YhdP are sufficient and required for membrane interaction.

Although these observations support a model in which the C-terminal helices of YhdP contact the OM, whether YhdP adopts an envelope-spanning architecture *in vivo* remains unclear. If true, interactions with the cell wall and OM would likely constrain the mobility of YhdP within the cell. To test this, *yhdP* was endogenously tagged with a HaloTag at the C-terminus, producing a strain that behaved the same as WT *E. coli* on SDS+EDTA agar plates, suggesting YhdP remained functional (Fig. S1B). The trajectories of the HaloTagged YhdP were tracked across 200 frames (Fig. S5A). These trajectories revealed little movement of YhdP within the cell (Fig. 2D). To quantify the mobility, the mean square displacement (MSD) was calculated, revealing immobile trajectories of YhdP (Fig. S5B) very similar to those of LptD and LptE, known to be immobile proteins at the OM^32^. The MSD was used to calculate the apparent diffusion coefficient of YhdP revealing that the majority of YhdP is immobile in cells (Fig. 2E). Of note, knocking out *dedD* did not alter the mobility of YhdP (Fig. 2E, S5C,D). In *E. coli*, IM proteins tend to have higher mobility than OMPs^33,32,34^. These data reveal that YhdP is as immobile as an OMP, supporting an envelope-spanning model where YhdP is constrained by the cell wall, and/or OM by its C-terminal helical loops.

### YhdP binds all major phospholipid classes

Having established YhdP as an envelope-spanning groove, the phospholipid-binding capabilities of YhdP were probed. Our previous *in vivo* crosslinking experiments captured phosphate-containing molecules within the groove^12^. To test if these molecules are likely phospholipids, the aforementioned YhdP_Cter_ΔH1&2 construct, which contains only a region of the hydrophobic groove and no membrane interacting sites, was tested for phospholipid binding by native MS. Native MS revealed YhdP_Cter_ΔH1&2 was primarily monomeric, with a minor dimer population (Fig. 3A). No bound phospholipids were detected in either population. Next, YhdP_Cter_ΔH1&2 was mixed with all three major classes of phospholipids in *E. coli*, namely dioleoyl phosphatidylethanolamine (DOPE), dioleoyl phosphatidylglycerol (DOPG) and cardiolipin (CDL 18:1), as well as an *E. coli* total lipid extract (Fig. 3B). From the spectra, most of the protein was observed in the *apo* state, but a subset of lipid-bound proteins were observed for all classes of phospholipid (Fig. 3B). A proportion of these contained multiple lipids, suggesting the YhdP_Cter_ΔH1&2 fragment can accommodate several lipids. Under the assay conditions, the YhdP fragment was shown to bind to a maximum of six CDL, three DOPE, or three DOPG (Fig. 3B). This stronger binding to CDL was likely due to its four hydrophobic tails. There was no obvious preference between lipid species, indicating that binding to the YhdP groove is not headgroup dependent, and rather driven by hydrophobicity of the acyl chains. Overall, these data suggest that the groove of YhdP can interact with all major classes of phospholipids in *E. coli*, and that phosphate-containing molecules previously captured *in vivo*^12^ along the length of the groove are likely phospholipids.

**Figure 3.**
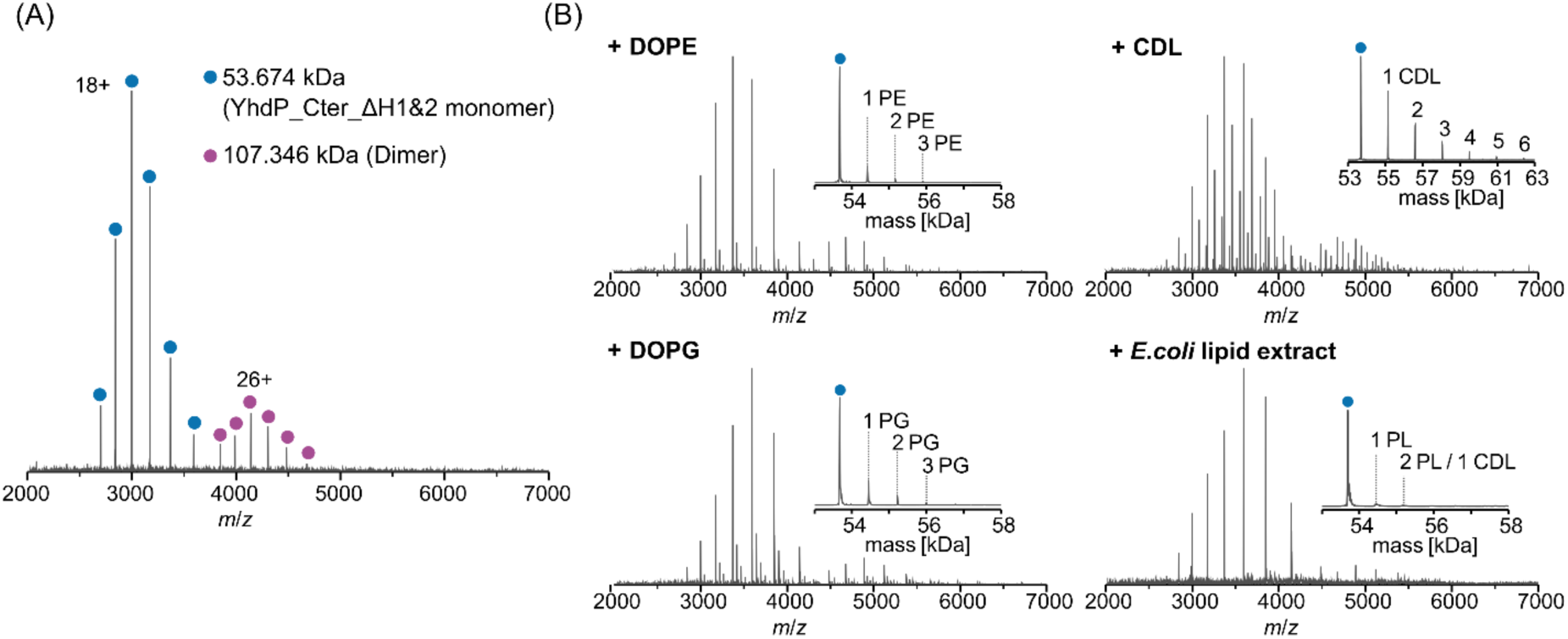
YhdP can bind to all classes of *E. coli* phospholipid. The purified YhdP_Cter_ΔH1&2 on its own (A) or with different phospholipids (B) were sprayed by native MS, with representative spectra shown. (A) The protein itself sprays mainly as monomers, in its *apo* state (blue dots). (B) After incubating with different phospholipids, the protein exists in its *apo* state, and different lipid-bound states. The deconvolution of the entire spectrum is shown as inlets.

### The disordered region of DedD is required to maintain the YhdP-DedD complex

Having established DedD as a native interactor of YhdP, the molecular details of this interaction was probed further. The AlphaFold model of the N-terminal section (residues 1-605) of YhdP complexed with full-length DedD was simulated in a membrane environment using coarse grained molecular dynamics (Fig. S6A). During the simulation, the interaction was maintained, with DedD forming a range of salt bridges with YhdP (Fig. S6A-C). DedD is composed of three domains, the TMH, the IDR and the SPOR domain (Fig. 4A). The salt bridges were formed between residues of the periplasmic domain of YhdP and either 1) the N terminal region (residues 1-86) of the DedD IDR, or 2) the SPOR domain of DedD (Fig. S6A-C). The overall architecture of the YhdP-DedD complex was consistent between AlphaFold and the MD simulations, except some of the salt bridges were predicted to form between different residues (Fig. S6A-D). Combining the top hits from both MD simulations and AF model, 14 residues on YhdP (Fig. S6E) were mutated to alanine individually to investigate whether altering any of the predicted salt bridges would disrupt the YhdP-DedD interaction. However, none of these mutations substantially impaired the expression and function of YhdP, nor altered its interaction with DedD (Fig. S7,8). Further, mutating these residues in clusters based on their locations (Fig. S6E), rendered YhdP unstable (Fig. S9) and therefore these mutants could not be probed further.

**Figure 4.**
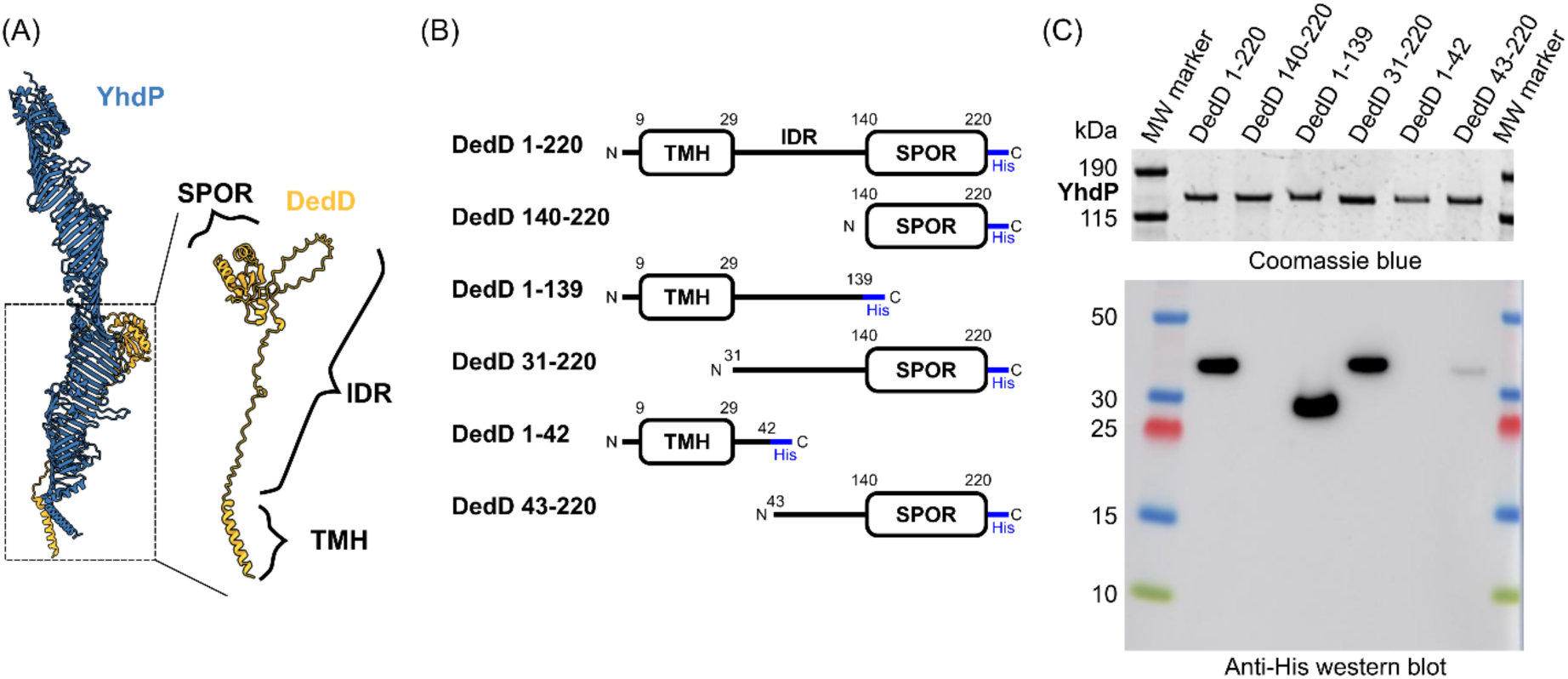
Mutagenesis of DedD to probe key interacting regions with YhdP. (A) Schematic diagram to show the domain architecture of DedD in the AlphaFold predicted YhdP-DedD model. The three domains of DedD are labelled. (B) The different constructs designed to test key regions of DedD for YhdP interaction. (C) Affinity purification of StrepII tagged WT YhdP co-expressed alongside His-tagged WT and mutant DedD constructs. The purification eluates were analysed by both Coomassie blue confirming purified YhdP, and anti-His western blot against DedD. Uncropped gels can be found in Fig. S10.

Instead, different truncated constructs of DedD were explored to determine the key domains of DedD that are required for YhdP interaction (Fig. 4B). These His-tagged DedD constructs were (1) full-length DedD (WT, residues 1-220); (2) the DedD SPOR domain only (140-220); (3) the TMH and IDR (1-139); (4) the IDR and SPOR domain (31-220); (5) the TMH with the N-terminus of the IDR (1-42); and (6) the C-terminus of the IDR followed by the SPOR domain (43-220). These DedD mutants were co-expressed with StrepII-tagged YhdP, followed by pull-down via YhdP. All DedD mutants expressed, as determined by western blot of whole cell lysates, though the DedD SPOR domain only (140-220) and the TMH with the N terminus of the IDR (1-42) were present at lower levels (Fig. S10A,B). Following YhdP purification, the TMH and IDR (1-139), and the IDR and SPOR (31-220) DedD constructs were pulled down at similar abundances to WT DedD (Fig. 4C, Fig. S10C,D). This indicates that neither the SPOR or TMH domains are essential for interaction with YhdP, and that the IDR may be the key interacting region. Similarly, DedD 140-220 (SPOR domain only) was not pulled down via YhdP, suggesting that the SPOR domain alone is not sufficient for YhdP interaction. Furthermore, there was little to no pull down of the DedD TMH with the N terminus of the IDR (1-42), or the C-terminus of the IDR followed by the SPOR domain (43-220). Importantly, it was confirmed that WT DedD did not non-specifically bind to the Strep resin (Fig. S10E), validating that pull-down of DedD was indeed due to interactions with YhdP. Together with the MD simulations and AlphaFold prediction these data point towards the DedD IDR being important in the YhdP-DedD interaction.

### DedD does not determine the cellular localisation of YhdP

While the molecular details of the YhdP-DedD interaction are now better understood, the role of DedD on YhdP function remains unknown. Previous studies of DedD have centred around its localisation at the septum during cell division, when expressed from a plasmid^20–22^. This is due to the SPOR domain interacting with denuded peptidoglycan, which is generated by amidases at the septum^20–22,26^. To investigate whether YhdP follows the same localisation pattern as DedD, the native cellular localisation of YhdP was probed.

First, to ensure that DedD is indeed septally localised at native levels, a C-terminally endogenously HaloTagged DedD strain was constructed and imaged (Fig. S5E). In dividing cells, and in agreement with the literature, DedD accumulated at the septum, but was also enriched at the cell poles (Fig. 5A). In non-dividing cells, DedD became mainly prevalent at the poles, with some localised along the cell length in a minority of cells. In contrast to DedD, native HaloTagged YhdP was scattered along the length of the cell, with no obvious enrichment of YhdP at the septum in dividing cells (Fig. 5B). Further, to determine if the absence of DedD altered YhdP localisation, the location of YhdP-HaloTag was tested in a *ΔdedD* strain. In this background, the location of YhdP was unaltered (Fig. S5F).

**Figure 5.**
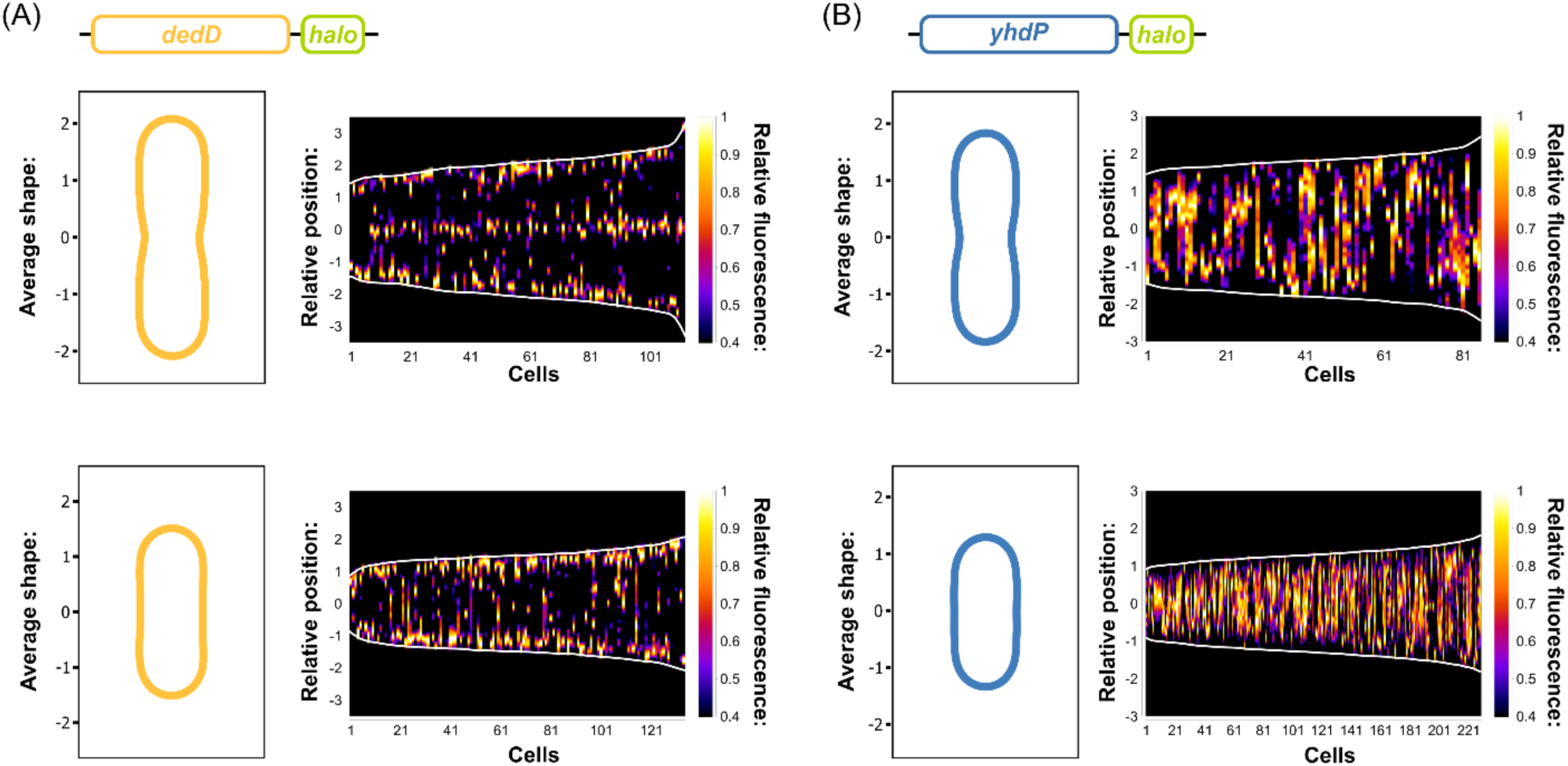
Super-resolution light microscopy of endogenous YhdP and DedD reveal they do not substantially co-localise. (A, B) Fluorescence profiles, depicted as demographs, of C-terminally Halo tagged DedD (A), and C-terminally Halo tagged YhdP (B). Averaged cell shapes for corresponding demographs are also shown. Data are split between dividing (upper) and non-dividing (lower) cells.

Together, these data reveal that native YhdP does not accumulate at any particular cellular localisation, native DedD is found at the septum and the poles, and that deletion of DedD does not alter YhdP location. Therefore it is unlikely that the primary role of the DedD-YhdP interaction is to localise YhdP. However, given the broad distribution of YhdP across the cell, it is likely that a proportion of YhdP interacts with DedD *in vivo*. Yet, the role of this DedD-YhdP interaction still remains unknown.

### DedD enriches phosphatidylethanolamine at the YhdP entrance

As DedD does not seem to be important for YhdP localisation at the septum, we hypothesised that it may instead be involved in facilitating YhdP function at the IM. To test whether DedD altered the membrane environment at the N-terminal entry of the YhdP groove (henceforward named the YhdP entrance), three coarse-grained MD simulations of full-length YhdP with or without DedD were carried out in a double-membrane system (Fig. 6A). From the same trajectory, lipid behaviours at the YhdP entrance were compared with the bulk membrane at increasing distances (6-18 Å, 18-30 Å, 30-42 Å). Lipid mobility decreased progressively towards the YhdP entrance in both the presence and absence of DedD, with DedD modestly decreasing these effects (Fig. S11A). The membrane also became increasingly tilted, relative to the bulk membrane, towards the YhdP entrance, with the largest increase within 6–18 Å in both conditions (Fig. S11B). Together, these changes showed that YhdP generated a locally constrained and deformed membrane environment, irrespective of the presence of DedD.

**Figure 6.**
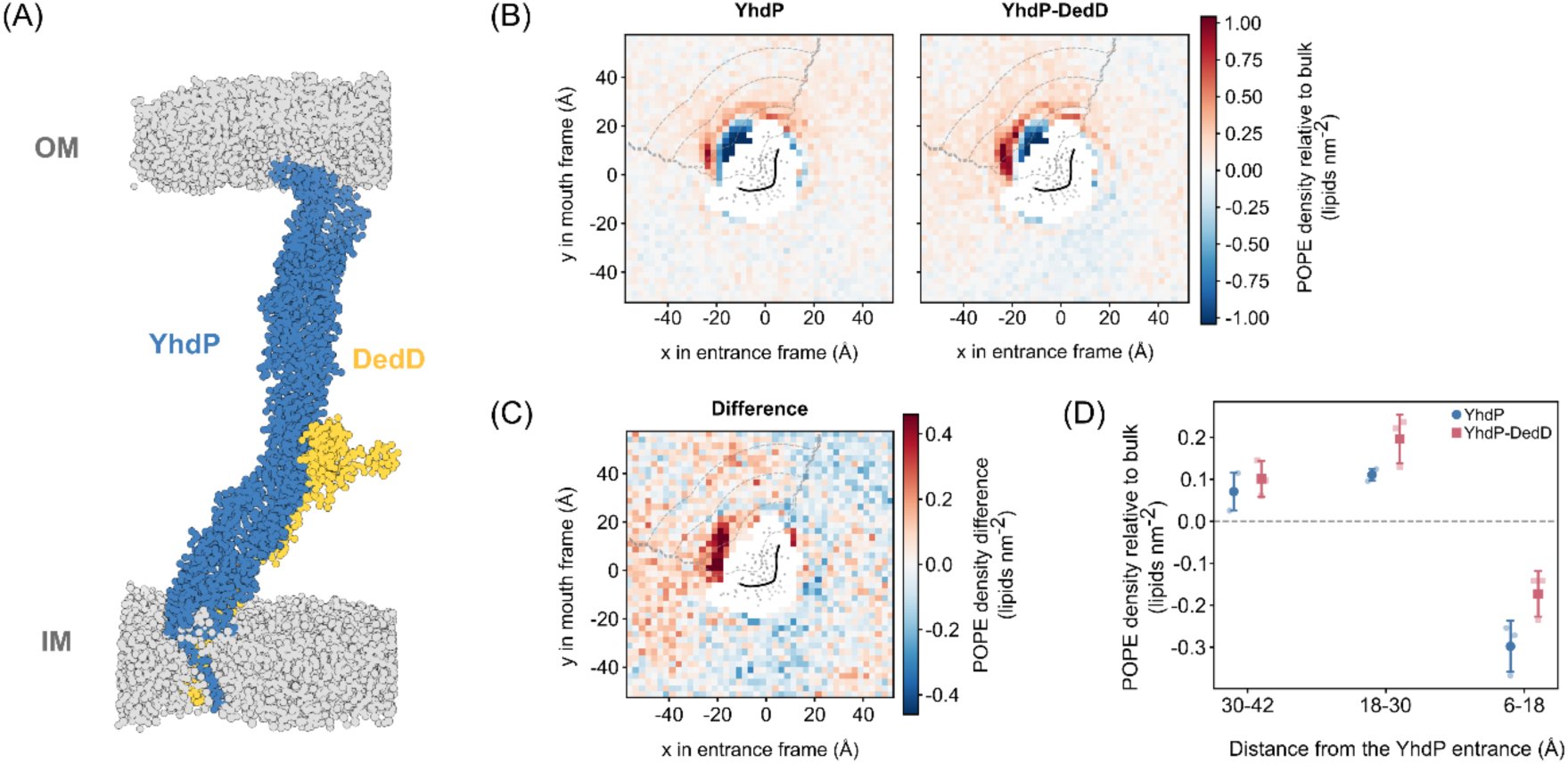
Molecular dynamics simulations indicate increased relative packing of phosphatidylethanolamine at the entrance of YhdP in the presence of DedD. (A) Representative snapshot of full-length YhdP–DedD simulated in a double-membrane system, showing YhdP spanning the inner (IM) and outer (OM) membranes and DedD associated with the IM region of YhdP. (B) Bulk-referenced POPE density maps around the YhdP entry for YhdP only (left) and YhdP–DedD (right) simulations. Red and blue indicate POPE density above and below the trajectory-specific bulk-membrane density, respectively. (C) Difference between the YhdP–DedD and YhdP only POPE density maps shown in (B). Positive values indicate increased POPE density in the presence of DedD. (D) POPE density relative to bulk within regions 30–42, 18–30 and 6–18 Å from the YhdP entry. Small points show the three independent trajectories and large points with error bars show the mean ± SD across trajectories. YhdP only is shown in blue and YhdP–DedD in red.

Next, the spatial distribution of phospholipids around the entrance of YhdP was analysed. Two density maps were constructed, either in the presence or absence of DedD. In both maps, POPE was not uniformly localised around YhdP, with altered density towards the YhdP entrance relative to the bulk membrane (Fig. 6B). Comparing both density maps, the presence of DedD enhances the POPE accumulation at the YhdP entrance (Fig. 6C). This effect was studied by quantifying POPE density as a function of distance from the YhdP entrance. Within 30-42 Å, POPE was enriched relative to bulk in both conditions, but DedD did not increase relative amounts of POPE (Fig. 6D). In contrast, within 18–30 Å, POPE was still enriched relative to the bulk membrane but this enrichment was greater in the presence of DedD. Within 6–18 Å, POPE was depleted relative to bulk in both conditions, but DedD substantially reduced this depletion. Neither POPG nor CDL showed an equivalent increase in density in the presence of DedD (Fig. S11C,D).

Together, these simulations showed that YhdP generated a local membrane environment characterised by reduced lipid mobility and increased membrane tilt. DedD modified this pre-existing YhdP-associated environment modestly in terms of mobility and tilt but consistently increased relative POPE density at the YhdP entrance.

## Discussion

The delivery of phospholipids from the inner membrane (IM) to the outer membrane (OM) is required for expansion of the Gram-negative bacterial cell envelope during growth. YhdP is the leading candidate for phospholipid transfer to the OM, yet the molecular mechanism has remained poorly understood. The cryo-EM data presented here reveal that YhdP contains a continuous cavity along its length, consistent with AlphaFold predictions^27^.

Endogenous YhdP is highly immobile within cells, with its C-terminal helical loops sufficient for membrane association *in vitro*, consistent with YhdP as an envelope-spanning bridge. The ability of YhdP to accommodate all three major classes of *E. coli* phospholipid further supports its proposed role in phospholipid transfer. The lack of an obvious headgroup preference suggests that hydrophobic interactions with lipid acyl chains contribute substantially to binding, allowing YhdP to accommodate the heterogeneous phospholipid composition of the *E. coli* membranes. This is supported by a recently published liposome transfer study of YhdP, where YhdP is able to transfer NBD tail-labeled PE and PG^13^. Together, these findings cement a model in which YhdP connects the IM and OM, positioning its hydrophobic groove across the periplasm to shield phospholipids during transfer.

The identification of DedD in this study and independently by Tan *et al*.^19^ suggests a functional interaction between the two proteins. The intrinsically disordered region (IDR) of DedD is required for interaction with YhdP, while neither the transmembrane nor SPOR domains are sufficient for binding. Disordered regions of this kind are a feature of other protein-protein interactions at the division site. FtsN, another SPOR domain-containing protein in *E. coli*, interacts with the septal peptidoglycan synthesis complex FtsWIQLB complex, via a short partially-formed periplasmic helix, to coordinate septal peptidoglycan synthesis^25,35,20,36^.

Under the conditions tested, deletion of *dedD* did not detectably alter the bulk localisation or mobility of YhdP, arguing against a straightforward role in recruiting YhdP to the division site. Instead, simulations indicate that DedD modestly increases PE occupancy around the YhdP IM entry point, raising the possibility that this interaction promotes phospholipid loading into the groove. Whether this reflects a direct role for DedD in lipid transfer or instead an indirect effect on the local membrane environment remains to be determined. It also remains to be established whether this effect is specific to DedD or arises more generally from the presence of an additional transmembrane helix adjacent to the YhdP entrance.

Several fundamental questions about YhdP-mediated phospholipid transfer remain unresolved. Although the ability of YhdP to bind multiple phospholipid classes is consistent with a phospholipid transfer role, it remains unclear how phospholipids are initially extracted from the IM and loaded into the groove, nor how they are inserted into the OM at the other end. In addition, like for the eukaryotic bridge-like lipid transfer proteins^37^, the preferred model is a diffusive process^39^. However, it is still unclear whether an energy source from the IM would aid transfer through YhdP.

Together, these findings provide biochemical information about how YhdP functions as an envelope-spanning phospholipid transfer bridge to aid in maintaining the OM of Gram-negative bacteria.

## Methods

### Antibiotic concentrations

In this study, carbenicillin, kanamycin and streptomycin were always used at 100, 50 and 100 µg/mL, respectively.

### Chromosomal modifications and endogenous purification of YhdP

Knockout of *yhdP* in *E. coli* BW25113 was performed using lambda red recombination with primers 1 and 2 (Table S1)^23^. An 11 amino acid (aa) linker (sequence: SAGSAAGSGEF) and Twin-Strep-tag, followed by a kanamycin resistant gene cassette was inserted at the N-terminus or C-terminus of *E. coli* BW25113 *yhdP*, using lambda red recombination with primers 3 - 9 (Table S1)^23,24^. For Halo-tagged, an 11 amino acid (aa) linker (sequence: SAGSAAGSGEF) and HaloTag, followed by a kanamycin resistant gene cassette was inserted at the C-terminus of *E. coli* BW25113 *yhdP* or *dedD*, using lambda red recombination with primers 6 - 13 (Table S1)^23,24^. The modified gene locus was transduced into *E. coli* BW25113 strain using P1 phage transduction^39^. The kanamycin cassette was removed using the pCP20 plasmid^40^. Colony PCR was performed to amplify regions flanking *yhdP* or *dedD*, to confirm the correct insertion by Sanger sequencing (SourceBiosciences).

To purify endogenous C-terminally Twin-Strep tagged YhdP from the engineered YCTS strain for single-particle cryo-EM, an overnight culture was grown from a single colony. Overnight culture was diluted 1:50 into 24 litres of Luria Broth (Miller’s) and grown at 37 °C with shaking to an OD600 of 0.80. Cells were harvested by centrifugation (4,000 x g, 20 min, 4 °C). Pellets were resuspended in Buffer A (50 mM HEPES, pH 7.5, 300 mM NaCl, 10% glycerol), supplemented with PMSF at a final concentration of 1 mM and EDTA-free protease inhibitor cocktail (Roche). Cells were lysed via an Emulsiflex C3 cell disruptor (Avestin). Cell lysates were clarified by centrifugation (16,000 x g, 30 min, 4 °C). Membrane pellets were collected by centrifuging the supernatant at 158,420 x g for 1 hr at 4 °C. Membrane pellets were solubilised in the Buffer A, with 1.2% 1.2% n-dodecyl β-D-maltoside (DDM) at 4 °C for 2 hours. The solubilised membrane was clarified by centrifugation (126,336 x g, 30 min, 4 °C). The supernatant was filtered through a 0.22 μm filter and incubated with Strep-Tactin®XT 4Flow® high-capacity resin (IBA Lifesciences) at 4 °C for 2 hours. The resin was washed with 100 column volumes (CV) of Buffer A, with 0.025% DDM. Bound proteins of interest were eluted in 10 CV of Buffer A, with 0.025% DDM and 50 mM biotin (IBA Lifesciences). Eluted protein was concentrated using a 100 kDa MWCO centrifugal filter (Sartorius). Concentrated protein was dialysed with a 3 kDa MWCO membrane (ThermoFisher) in 25 mM HEPES, pH 7.5, 150 mM NaCl buffer overnight at 4 °C for Cryo-EM (see “**Single-particle cryo-EM sample preparation and data collection**”).

To prepare samples for proteomics experiments, overnight cultures of both wild-type *E. coli* BW25113 (WT) and the C-terminally Twin-Strep tagged YhdP strain were grown from single colonies. Overnight cultures were diluted 1:50 in Luria Broth (Miller’s) re-grown at 37 °C with shaking to OD600 of 0.80 (4 litres for each strain, in triplicate). Cells were harvested by centrifugation (4,000 x g, 20 min, 4 °C). Cell weight masses were normalised. Pellets were resuspended in Buffer C (20 mM HEPES, pH 7.5, 150 mM NaCl), supplemented with PMSF at a final concentration of 1 mM. Cells were lysed through sonication (7s on, 3s off, for 90s in total). Cell lysates were clarified by centrifugation (16,000 x g, 30 min, 4 °C). Membrane pellets were collected by centrifuging the supernatant at 158,420 x g for 1 hr at 4 °C. Membrane pellets were frozen and stored at −70 °C, until purification. Membrane pellets were solubilised in Buffer C, with 1.2% DDM at 4 °C for 2 hours. The solubilised membrane was clarified by centrifugation (126,336 x g, 30 min, 4 °C). Supernatant was filtered through a 0.22 μm membrane and incubated with 150 μl CV Strep-Tactin®XT 4Flow® high-capacity resin (IBA Lifesciences) at 4 °C for 2 hours. The resin was washed with 50 CV of Buffer C, with 0.025% DDM. 50 μl of resin was washed in PBS twice and stored in fresh 8 M urea 100 mM ammonium bicarbonate buffer at −70 °C, until sample preparation for proteomics.

### Single-particle cryo-EM sample preparation and data collection

3 μl of freshly purified endogenous YhdP sample, at 0.5 mg/ml, was applied on to glow-discharged R 1.2/1.3 Cu 300 mesh grids with 2 nm continuous carbon (Quantifoil). The grid was glow-discharged for 15 s at 15 mA using EM ACE200 (Leica). Vitrification was performed using Vitrobot Mark IV (ThermoFisher Scientific), at 4°C and 100% humidity, for 3 seconds at a blot force of 3, before being plunged into liquid ethane. Cryo-EM data were acquired on a 300 kV Titan Krios G3 (FEI) with a K3 direct electron detector (Gatan) and BioQuantum imaging filter. A total of 26,488 movies were acquired automatically in counted super resolution mode using EPU software (Thermo Fisher Scientific) at 105,000x magnification, corresponding to a calibrated pixel size of 0.835 Å. Movies were collected with a target defocus range of −3.0 to −1.4 µm at a total dose of 40.1 e-/Å2 fractionated into 40 frames.

### Cryo-EM data processing

Movies were imported into CryoSPARC (v4.7.1) for all processing. The processing was summarised in Figure S2. Beam-induced motion and contrast transfer function (CTF) were corrected by patch-based motion correction and patch-based CTF estimation, respectively. Micrographs with too thin ice and broken carbon support layers were discarded. The resulting 21,222 micrographs were randomly split into a batch size of 5,000. One batch was used for picking parameter optimisation. Blob picking, with the elliptical blob, was used for initial particle picking, followed by 2D classification to generate a good template to train Topaz ^41^. The trained Topaz model was used to pick particles from the entire dataset. Iterations of 2D classifications were used to clean up the particle set. Classes with relatively well aligned particles were selected. An initial 3D volume was generated using Ab-Initio reconstruction and non-uniform refinement, with default settings. The particle set was cleaned up in 3D using heterogeneous refinement, with two decoy volumes. A range of jobs, such as 3D classification with or without focused mask at different filter resolutions, and local refinement with various masks, were tried to improve the resolution of this endogenous YhdP sample. However, no confident secondary structure could be visualised in the density map. The particle set was further cleaned up, following the HR-HAIR pipeline^42^. 2D classes with the best alignment were used for Ab-Initio reconstruction with HR-HAIR parameters. The final 3D volume at 8.30 Å was generated using non-uniform refinement.

### On-bead digestion and peptide identification by LC-MS/MS

Resin with proteins of interest bound stored in 8 M urea 100 mM ammonium bicarbonate were reduced with Tris(2-carboxyethyl) phosphine (TCEP) and alkylated with 2-chloroacetamide. Samples were digested with 1 µg of LysC for 2 hr at 37 °C. The urea concentrations of the samples were diluted to 2 M, and calcium chloride was added to a final concentration of 2 mM. Peptides were digested overnight with 100 ng trypsin at 37 °C. Digestion was stopped with formic acid, at a final concentration of 5%. Supernatants were collected after centrifugation (14,000 rpm, 30 min, 4 °C). C18 resin was used to desalt the digested peptide solution.

Peptides were separated by nano liquid chromatography (Thermo Scientific Ultimate 3000) coupled to a Q Exactive mass spectrometer equipped with an Easy-Spray source (Thermo Fisher Scientific). Peptides were trapped onto a C18 PepMap100 precolumn (300 µm i.d., 5 mm length, 100 Å, Thermo Fisher Scientific) using Solvent A (0.1 % formic acid, HPLC grade water). The peptides were further separated onto an Easy-Spray C18 column (75 µm i.d., 50 cm length, Thermo Fisher Scientific) using a 30 min linear gradient (15 % to 38 % solvent B (0.1 % formic acid in acetonitrile)) at a flow rate 200 nL/min. The raw data were acquired on the mass spectrometer in a data-dependent acquisition mode (DDA). Full-scan mass spectra were acquired in the Orbitrap (Scan range 350-1500 m/z, resolution 70,000; AGC target 3e6, maximum injection time 50 ms). The 10 most intense peaks were selected for higher-energy collision dissociation (HCD) at 30 % of normalized collision energy. HCD spectra were acquired in the Orbitrap at resolution 17,500, AGC target 5e4, maximum injection time 120 ms with fixed mass at 180 m/z. Charge exclusion was selected for unassigned and 1+ ions. The dynamic exclusion was set to 10 s.

Tandem mass spectra were searched using Sequest HT in Proteome Discoverer software version 1.4, against an *Escherichia coli* K12 protein sequence database, downloaded from UniProt, containing 4,402 protein entries (Proteome ID: UP000000625) and 292 common contaminants. Carbamidomethylation on cysteine was set as a fixed modification; oxidation on methionine, N-terminal acetylation, deamidation on asparagine and glutamine and carbamylation of lysine were set as variable modifications. Two missed cleavages were permitted. Peptide mass tolerance was set at 20 ppm on the precursor and 0.6 Da on the fragment ions. Data was filtered at FDR below 1 % at the peptide level.

### Construction of plasmids for protein purification

For full-length YhdP and DedD co-expression, the gene locus of *yhdP* or *dedD* was amplified from *E. coli* BW25113 and cloned by Gibson assembly into a pCDFDuet or pET26b vector, respectively. YhdP was cloned with a StrepII tag at the C-terminus, resulting in plasmid pGLI196, and DedD with a C-terminal 10x His tag, resulting in plasmid pQT028. Both plasmids were co-transformed into T7 express competent *E. coli* cells (NEB) for co-expression.

For soluble YhdP expression, YhdP residues 1101 to 1266 were cloned by Gibson assembly into a pET26 vector, with a MBP at the N-terminus and a StrepII tag at the C-terminus, resulting plasmid pQT052 (YhdP_Cter_WT). For plasmid pQT059 (YhdP_Cter_ΔH1&2), YhdP residues 1121-1144 and 1203-1237 on pQT052 were mutated to ‘GS’ and ‘GSGS’ linkers, respectively, by Gibson assembly. Plasmids were transformed into T7 express competent *E. coli* cells (NEB) for expression.

### Protein over-expression and purification

For expression of all constructs in T7 express, overnight cultures were diluted 1:50 in Luria Broth (Miller’s), supplemented with appropriate antibiotics and grown at 37 °C with shaking to OD600 of 0.80. Cultures were cooled down for 1 hour at 4 °C and then induced with IPTG, at a final concentration of 1mM, at 15 °C with shaking overnight. Cells were harvested at 4,500 x g for 15 min at 4 °C.

For purification of the YhdP-DedD complex, cell pellets were resuspended in Buffer A (50 mM HEPES, pH 7.5, 300 mM NaCl, 10% glycerol), supplemented with PMSF at a final concentration of 1 mM. Cells were lysed via Emulsiflex C3 cell disruptor (Avestin). Cell lysates were clarified by centrifugation (16,000 x g, 30 min, 4 °C). Membrane pellets were collected by centrifuging the supernatant at 158,420 x g for 1 hr at 4 °C. Membrane pellets were solubilised in Buffer A, with DDM at 4 °C. The solubilised membrane was clarified by centrifugation (126,336 x g, 30 min, 4 °C). Supernatant was filtered through a 0.22 μm membrane and incubated with 1 ml column volume (CV) Strep-Tactin®XT 4Flow® high-capacity resin (IBA Lifesciences) at 4 °C for 2 hours. The resin was washed with 50 CV of Buffer A, with 0.025% DDM. Bound proteins of interest were eluted in 10 CV of Buffer A, with 0.025% DDM and 50 mM biotin (IBA Lifesciences). Eluted protein was incubated with 500 μl Ni Sepharose 6 Fast Flow resin at 4 °C for 1 hour. The resin was washed with 8 CV of Buffer A, with 0.025% DDM and 40 mM imidazole. Proteins of interest were eluted with 5 CV of Buffer A, with 0.025% DDM and 250 mM imidazole. The eluted complex was concentrated using a 100 kDa MWCO centrifugal filter (Sartorius). Concentrated protein was dialysed with a 3 kDa MWCO membrane (ThermoFisher) in 25 mM HEPES, pH 7.5, 150 mM NaCl buffer overnight at 4 °C.

For purification of soluble YhdP constructs, cell pellets were resuspended in Buffer B (50 mM HEPES, pH 7.5, 500 mM NaCl, 10% glycerol), supplemented with PMSF at a final concentration of 1 mM. Cells were lysed via Emulsiflex C3 cell disruptor (Avestin). Cell lysates were clarified by centrifugation (36,000 x g, 1 hr, 4 °C). Supernatant was filtered through a 0.45 μm membrane and incubated with 500 μl CV Strep-Tactin®XT 4Flow® high-capacity resin (IBA Lifesciences) at 4 °C for 2 hours. The resin was washed with 100 CV of Buffer B. Bound proteins of interest were eluted in 10 CV of Buffer B, with 50 mM biotin (IBA Lifesciences). The eluted complex was concentrated using a 30 kDa MWCO centrifugal filter (Sartorius). Concentrated protein was loaded onto Superose 6 increase 10/300 GL column (Cytiva) or Superdex 200 increase 10/300 GL column (Cytiva), equilibrated with 20 mM HEPES, pH 7.5, 150 mM NaCl. Fractions across the peak were analysed by SDS-PAGE. Fractions, containing proteins of interest, were pooled and concentrated using a 30 kDa MWCO centrifugal filter (Sartorius). Proteins were flash frozen in liquid N2 and stored at −70 °C, until use.

### Molecular dynamics system set-up, simulations and analysis

All simulations were performed within a charge neutralised system with the addition of ions corresponding to a solution with a 0.15 M NaCl concentration. Full-length YhdP, with or without DedD, were coarse-grained using martinize2^43^ with the Martini2.2 force field^44^ and an elastic network with a force constant of 500 kJ mol⁻¹ nm⁻². Double-membrane systems were assembled using COBY^45^ with each replica given a different random seed. The inner membrane was composed of POPE:POPG:CDL2 at a 15:4:1 ratio in both leaflets. The outer membrane contained rough LPS in the external leaflet and POPE:POPG:CDL2 at an 18:1:1 ratio in the periplasmic leaflet. A modified FhuA nanopore was included within the double-membrane system to enable exchange of coarse-grained water between the aqueous compartments, as described previously^12^. Pilot fragment simulations containing YhdP residues 1–605 together with DedD were set up using the same procedure described above, in a model inner membrane composed of POPE:POPG:CDL2 at a 15:4:1 ratio in both leaflets.

The velocity rescale (V-rescale) thermostat^46^ with a coupling constant of 1 ps was used to maintain the system temperature at 323 K. Pressure was maintained at 1 bar using the c-rescale barostat^47^ with a coupling constant of 4 ps and semi-isotropic pressure coupling. All simulations using Martini2.2 used reaction-field electrostatics with a relative dielectric constant of 15 and electrostatic and van der Waals cut-offs of 1.1 nm. The full systems were first energy minimised using the steepest descent algorithm. Equilibration consisted of 0.5 ns within the NVT ensemble using a 2 fs integration time step, followed by successive NPT simulations of 1, 10 and 20 ns using integration time steps of 5, 10 and 12.5 fs, respectively. Restraints on the protein backbone and lipid headgroups were progressively reduced during equilibration. During the final equilibration stage and production simulations, a flat-bottom restraint was applied in the x dimension between the YhdP C-terminal region and the FhuA nanopore, maintaining their separation between 5 and 10 nm. Three independent simulations of 2.5 μs were carried out for YhdP alone and for YhdP in the presence of DedD, using a 12.5 fs integration time step. The pilot YhdP–DedD fragment systems were equilibrated using the same general procedure, with an initial 1 ns simulation within the NVT ensemble followed by successive NPT simulations of 5 and 20 ns. The integration time step was increased from 2 to 10 and then 20 fs, while restraints on the protein backbone and lipid headgroups were progressively reduced. Production simulations used a 20 fs integration time step.

Analysis was performed using MDAnalysis^48,49^, MDTraj^50^ and in-house Python scripts. Following assessment of bulk-membrane equilibration, the final 1 μs of each trajectory (1.5–2.5 μs) was used for analysis. The upper inner-membrane leaflet adjacent to the N-terminal mouth of YhdP was analysed. A single mouth definition was used for all replicas and both conditions, defined by a curve through YhdP backbone beads at residues 31, 60, 61, 63, 95 and 97. Lipids were grouped into regions 6–18, 18–30 and 30–42 Å from this boundary. Bulk membrane was defined independently within each trajectory as the region 70–100 Å from the protein using three-dimensional minimum-image distances. Lipid mobility was calculated from identity-tracked lipid displacements projected into the local membrane tangent plane. Local membrane tilt was calculated from fitted leaflet surfaces and defined relative to the instantaneous global inner-membrane normal. Temporal subdivisions of trajectories were used only as robustness checks.

### Construction of *yhdP* and *dedD* mutants for mutagenesis experiments

For YhdP mutants, primers were designed to introduce mutations in desired residues to alanine (codon GCG). PCR products were inserted into a pET26b backbone (by XbaI and NheI), by Gibson assembly. Resulting plasmids were isolated and transformed into competent *E.coli* BW25113 *ΔyhdP* for phenotypic assays. They were also co-transformed with pBM027 (pCDF-Duet harbouring WT His-tagged DedD) into competent *E. coli* T7 express cells, for pull-downs.

For DedD truncation mutants, different regions of *dedD* were amplified by PCR. A PelB signal sequence was introduced for DedD 140-216, 31-216 and 43-216 constructs. PCR products were inserted into pCDF-Duet backbone (digested by NcoI and XhoI), by Gibson assembly. Resulting plasmids were isolated and co-transformed with pQT033 (pET26b harbouring WT StrepII tagged YhdP into competent *E. coli* T7 express cells, for pull-downs.

### Screening expression of *yhdP* mutants by phenotypic assays and western blot

All YhdP mutants were tested in phenotypic assays with four *E. coli* BW25113 controls: WT, *ΔyhdP*, *ΔyhdP* with pQT033 (pET26b with WT StrepII-tagged YhdP), and *ΔyhdP* with empty pET26b. All mutant and control pET26b plasmids were transformed into chemically competent *E. coli* BW25113 *ΔyhdP*. Cells were grown overnight at 37°C, in cultures supplemented with appropriate antibiotics. 10-fold serial dilutions of the overnight cultures in 96-well plates were spotted on LB agar with or without 0.5% SDS + 0.55 mM EDTA. Plates were incubated overnight at 37°C. For all mutants, three biological replicates were performed.

Alongside YhdP mutant phenotypic assays, cell lysate samples from overnight cultures were boiled at 95 °C in SDS-Loading buffer and analysed via anti-Strep western blot. For western blots, gels were transferred to a nitrocellulose membrane. Membranes were blocked with 5% skimmed milk in PBS-T buffer, followed by overnight incubation with anti-Strep antibody (Abcam ab183907, a dilution ratio of 1:1,000). Membranes were washed in PBS-T buffer and incubated with the secondary antibody (Abcam ab6721, a dilution ratio of 1:5,000). After washing in PBS-T buffer, membranes were incubated with ECL substrate Working Solution (ThermoFisherScientific, 32106) and imaged by Hyperfilm ECL (Amersham).

### Small-scale pull-downs on StrepII-tagged YhdP

T7 express cells, with different *yhdP* and *dedD* plasmids, were grown and induced the same way as mentioned above, but in 10 ml cultures. Cell pellets were resuspended in PBS, with 0.5 mM EDTA, 1 mg/ml lysozyme and 1 μg/ml benzonase. Cells were lysed by eight cycles of freeze thaw, between liquid nitrogen and 37 °C in a heat block. Soluble materials were removed by centrifuging at 16,000 x g for 30 min at 4 °C. Pellets were solubilised in Buffer D (50 mM HEPES, pH 7.5, 150 mM NaCl, 10% glycerol), with 1.2% DDM. Insoluble materials are removed by centrifuging at 16,000 x g for 15 min at 4 °C. Supernatant was incubated with Strep-Tactin®XT 4Flow® high-capacity resin (IBA Lifesciences) at 4 °C. The resin was washed with 160 column volume (CV) of Buffer D, with 0.025% DDM. Bound proteins of interest were eluted in 6 CV of Buffer D, with 0.025% DDM and 50 mM biotin (IBA Lifesciences). Elutions are analysed by SDS-PAGE, followed by Coomassie-blue staining or anti-His western blot. Gels were transferred to a nitrocellulose membrane. Membranes were blocked with 5% skimmed milk in PBS-T buffer, followed by overnight incubation with anti-His antibody (Abcam ab18184, a dilution ratio of 1:2,000). Membranes were washed in PBS-T buffer and incubated with the secondary antibody (Abcam ab6789, a dilution ratio of 1:5,000). After washing in PBS-T buffer, membranes were incubated with ECL substrate Working Solution (ThermoFisherScientific, 32106) and imaged by Hyperfilm ECL (Amersham).

### Fluorescence microscopy and image analysis

Overnight cultures of bacterial strains in M9 media were set up from a single colony and grown at 37 °C. Overnight cultures were diluted 1:100 in M9 media and grown to OD600 0.2. Cells were harvested and resuspended in 100 μl M9 media. Janelia Fluor^®^ JFX554 ligand (Promega) was incubated with cell solution, at a final concentration of 150 nM, for 1 hr. Cells were pelleted and washed with new M9 media 10 times, to remove non-specific dyes. After 30 min recovery at 37 °C, cells were added onto the agarose pad (1% w/v), with SecureFrame. Cells were imaged using the Oxford Nanoimager-S with a 100×/1.49 oil immersion objective lens and a pixel size of 117 nm. Cells were first imaged with 560 nm excitation (fluorescence) for 200 frames, and 650 nm transillumination (brightfield) for 100 frames. Each frame had a 30 ms acquisition time. The 560 nm and 650 nm lasers were used at 20% and 10% power, respectively, for excitation.

Demographs were plotted using the MicrobeJ plug-in in Fiji (v.2.16.0/1.54p)^51,52^. All brightfield frames of each acquisition were averaged and transferred into a binary mask. All fluorescent frames of each acquisition were averaged as the representative fluorescent image. Cells were manually classified as dividing or non-dividing. Averaged cell shapes from each classification were plotted in MicrobeJ. Fluorescence profiles were plotted against mean fluorescence per pixel along the segmentation-determined cell length axis and were ranked by cell length.

Single-molecule tracking was performed using Trackmate (v.7.14.0) and MSDanalyser^53–55^. Fluorescent spots were detected using the DoG detector, with an estimated object diameter of 0.5 µm and a quality control of 5. Tracks were formed using the LAP tracker, with a maximum linking distance of 0.5 µm and a maximum frame gap of 2. Tracks with at least 20 spots were exported to MSDanalyser. MSD was calculated through MSDanalyser in MATLAB (version R2025a). The first 4 MSD points of each trajectory were used to fit a linear regression. Apparent diffusion coefficient is calculated with the anomalous diffusion exponent of 1.

### Liposome reconstitution and co-sedimentation experiments

*E. coli* total lipid extract (Avanti) was dissolved in chloroform. A lipid film was formed by drying the chloroform-solubilised lipids with nitrogen gas. A vacuum desiccator was used to remove residual chloroform overnight. The lipid film was rehydrated to 20 mg/ml in the liposome reconstitution buffer (100 mM HEPES, pH 7.5, 100 mM NaCl). To form liposomes, the lipid solution was extruded through 200 nm membrane filters (Merk Millipore) using a mini-extruder (Avanti) 31 times to form liposomes. Reconstituted liposomes were stored at 4°C until use.

Aggregates from purified 8 μM pQT052 (YhdP_Cter_WT) or pQT059 (YhdP_Cter_ΔH1&2) samples were removed by centrifuging at 100,000 x g for 30 min at 4 °C. Supernatants were incubated with 5 mg/ml liposome at room temperature for 2 hr. The first liposome pellet was collected by centrifuging at 100,000 xg for 30 min at 4°C. Supernatants were collected and the liposome pellet was washed with the liposome reconstitution buffer. The second liposome pellet was collected again by centrifuging at 100,000 x g for 30 min at 4 °C. Supernatants were collected and the pellet was resuspended in the liposome reconstitution buffer. Samples were collected at each step, with volume normalised, for SDS-PAGE. Three biological repeats were performed.

### Native mass spectrometry

The YhdP_Cter_ΔH1&2 mutant was buffer-exchanged into 200 mM ammonium acetate (pH 7.0) containing 2x critical micelle concentration (CMC) n-octyl-β-D-glucopyranoside (OG) using 40 kDa cutoff Zeba spin desalting columns (Thermo Fisher Scientific). The YhdP-DedD complex was buffer-exchanged into 200 mM ammonium acetate (pH 7.0) with 2xCMC n-decyl-β-maltoside (DM) using the same protocol. The protein solutions were loaded into borosilicate nano-emitters pulled in-house using a P97 Micropipette Puller (Sutter Instrument Corporation) and coated with gold by an Agar Auto Sputter Coater. Native mass spectra were recorded on a Q Exactive UHMR mass spectrometer (Thermo Fisher Scientific). Proteins were ionised by nano-electrospray ionisation by applying a voltage of 0.9–1.1 kV, and all spectra were measured in positive ion mode.

The native mass spectrum of the YhdP C-ter helices deletion mutant (ca. 5 µM) was recorded from *m*/*z* 2,000–10,000 with the following instrument parameters: capillary temperature: 200 °C; desolvation voltage: −250 V; collisional activation in the HCD cell: 100 V; trapping gas pressure setting: 5.0; Orbitrap resolution: 12,500 @ *m*/*z* 400. To investigate lipid binding, lipid solutions were prepared as follows: DOPG, DOPE, CDL (all 18:1), and *E. coli* lipid extract in chloroform (all purchased from Avanti) were dried in glass vials in a centrifugal evaporator overnight. The lipid film was resuspended to a stock concentration of 5 mM in 200 mM ammonium acetate (pH 7.0) with 2xCMC tetraethylene glycol monooctyl ether (C8E4). The stock solutions were diluted in 200 mM ammonium acetate (pH 7.0) with 2xCMC OG to a final concentration of 100 µM. These solutions were mixed 1:2 (v:v) with the buffer-exchanged YhdP_Cter_ΔH1&2 mutant, resulting in a final mixture of ca. 3 µM protein and 33 µM lipid. For the *E. coli* lipid extract, the total lipid concentration was roughly estimated assuming an average molecular weight of 750 g/mol. Each protein-lipid mixture was analysed in two technical replicates by native MS using the same settings as above and collisional activation in the HCD cell set to 200 V. The native mass spectrum of the YhdP-DedD complex was recorded from *m*/*z* 1,000–10,000 with the following instrument parameters: capillary temperature: 200 °C; desolvation voltage: −100 V; collisional activation in the HCD cell: 250 V; trapping gas pressure setting: 8.0; Orbitrap resolution: 25,000 @ *m*/*z* 400. All native mass spectra were deconvolved in UniDec with mass sampling every 1.0 Da to determine intact protein masses^56^.

### Structural prediction and visualisation

All AlphaFold models were generated using the AlphaFold server with AlphaFold 3^27^. All AlphaFold models and cryo-EM data were visualised and edited using ChimeraX (version 1.9)^57,58^.

## Acknowledgements

The authors thank Matthew Hankins and Harry Lee for providing feedback on the manuscript and Matthew Hankins for advice on liposome assay set-up. The authors thank Dr Marjorie Fournier and Dr Vaishnavi Ravikumar of The Advanced Proteomics Facility, Department of Biochemistry, University of Oxford, for their support and assistance with the proteomics experiments and data analysis. The authors thank the COSMIC cryo-EM facility (Department of Biochemistry, University of Oxford) for their support and assistance in this work.

---

**Supplementary Figure 1.**
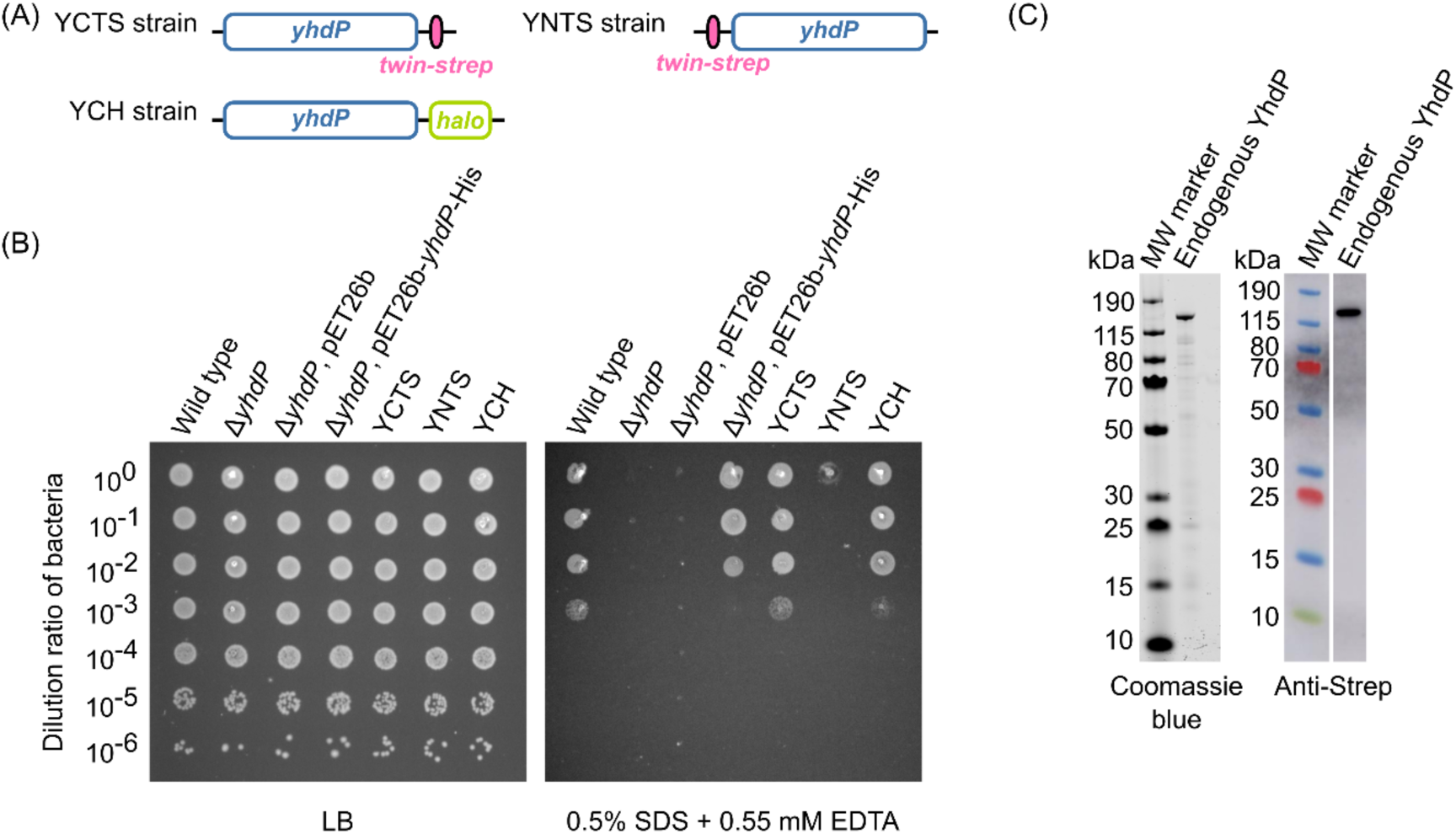
Phenotypic assay and purification of endogenously tagged strains. (A) Three endogenously tagged *yhdP* strains were constructed in this study. The gene schematic of the strains used are shown. (B) 10-fold dilution series of indicated strains were spotted on LB agar plates with or without SDS+EDTA. Representative plates are shown. (C) A Coomassie-blue stained gel and anti-Strep western blot indicate the successful purification of endogenous YhdP from the C-terminally Twin-Strep tagged strain.

**Supplementary Figure 2.**
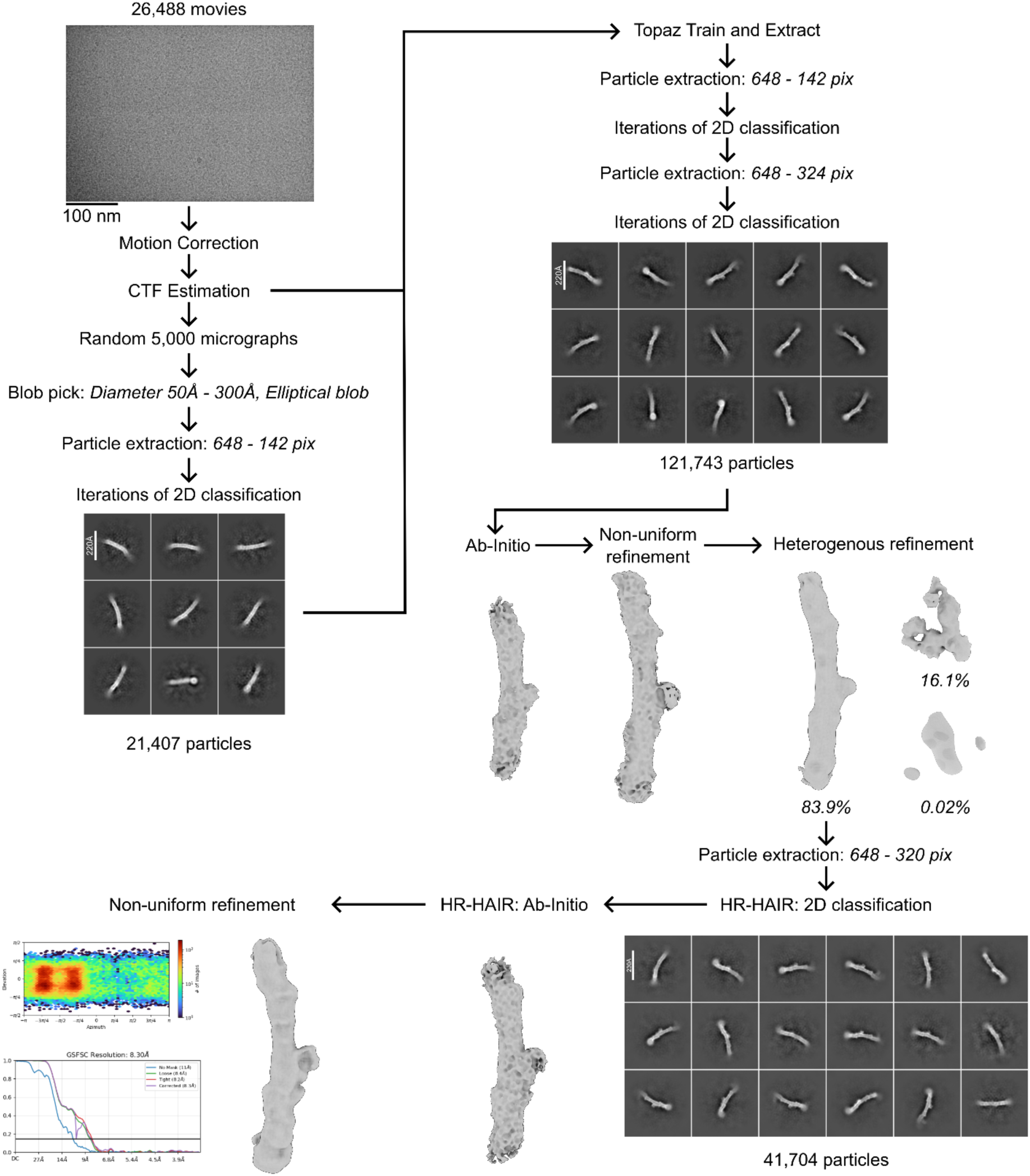
Single-particle cryo-EM processing pipeline of the endogenous YhdP sample. Data processing was performed in CryoSPARC.

**Supplementary Figure 3.**
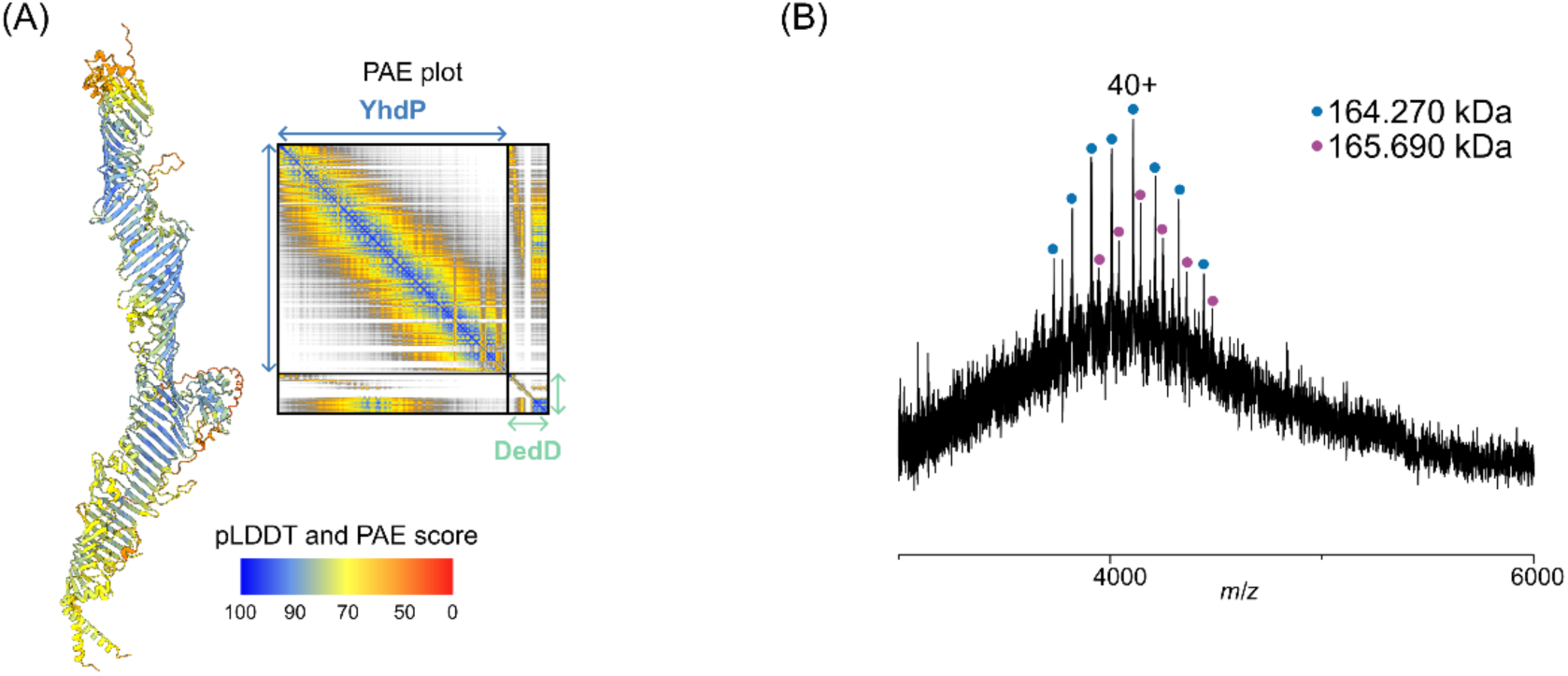
Biochemical and biophysical validation of the YhdP-DedD complex. (A) AlphaFold model of the YhdP-DedD complex is coloured by pLDDT scores. The PAE plot is shown as inlet. (B) Native MS spectrum of the YhdP-DedD complex confirms a 1:1 stoichiometry. The native MS spectrum revealed two species with molecular weights of 164,270 Da and 165,690 Da indicating a 1:1 stoichiometry (based on the theoretical molecular weight of a 1:1 complex of 164,403 Da). The decrease of 133 Da between the theoretical mass and the 164,270 Da peak could correspond to the loss of methionine during protein expression. The 165,690 Da peak, which is heavier than the first peak by 1,420 Da, could indicate binding of a small molecule, such as cardiolipin. However, the spectrum was not at high enough resolution to identify this adduct unambiguously.

**Supplementary Figure 4.**
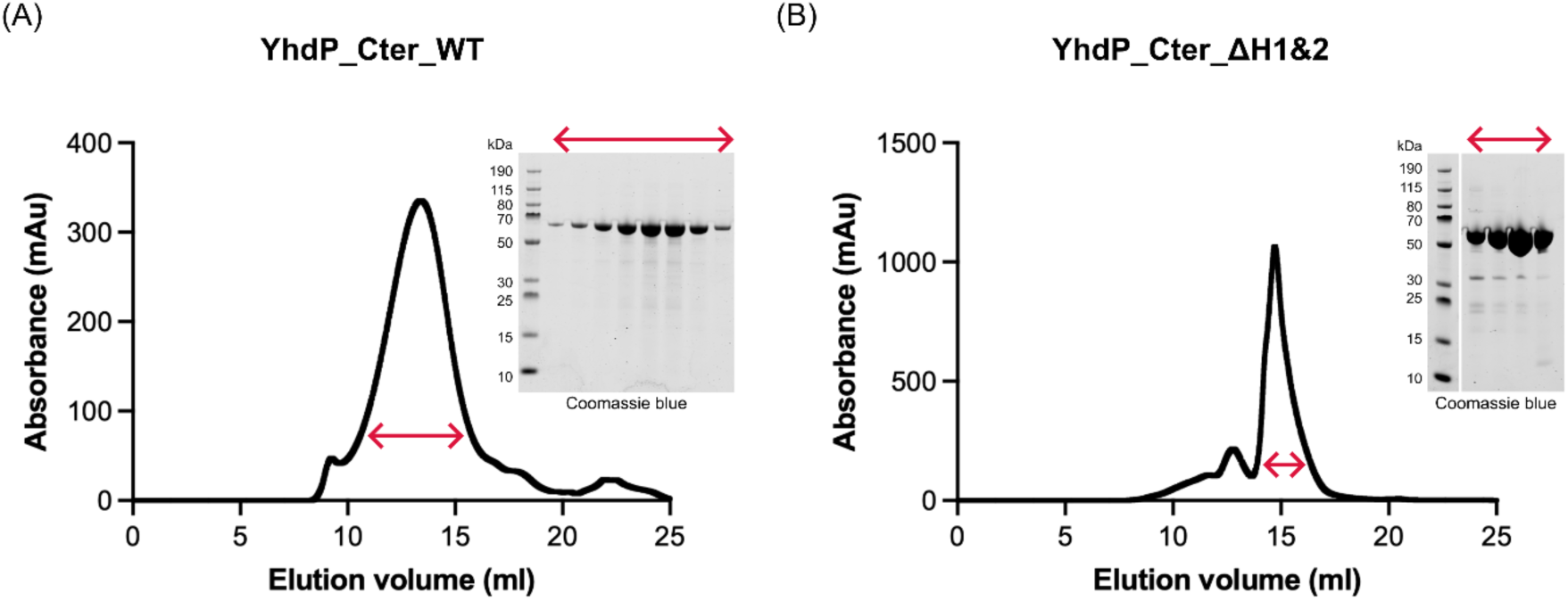
Purification of YhdP_Cter_WT and YhdP_Cter_ ΔH1&2. Both YhdP_Cter_WT (A) and YhdP_Cter_ ΔH1&2 (B) constructs were purified as described in Methods. Size exclusion chromatography traces are shown here, with samples from the main peaks shown on Coomassie-blue stained gels.

**Supplementary Figure 5.**
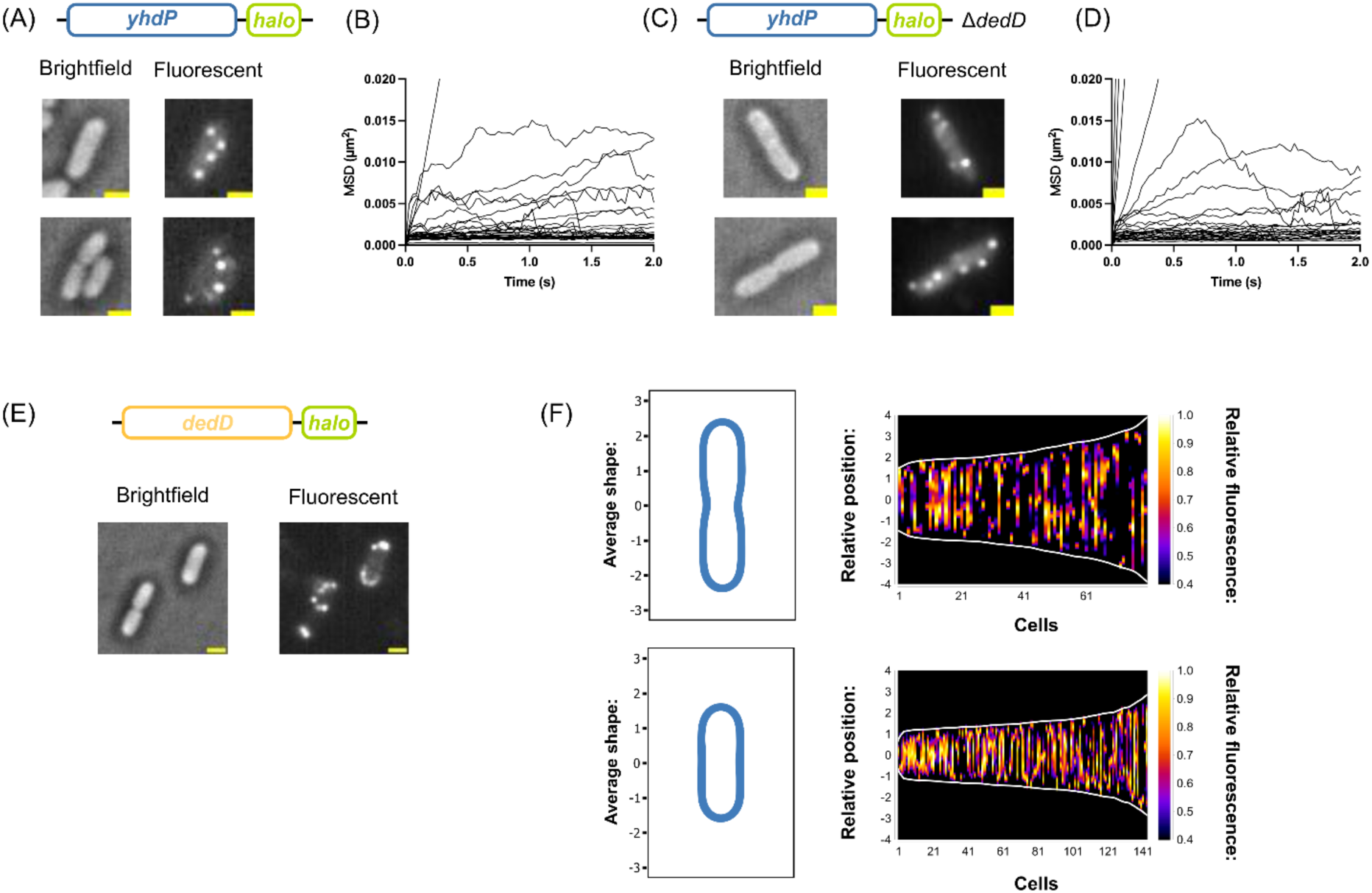
Fluorescent microscopy of YhdP and DedD *in vivo* at endogenous levels. (A) Representative brightfield and fluorescent images are shown for C-terminally Halo tagged YhdP. These images are generated by averaging all frames. Scale bar (yellow): 1 µm. (B) Computed mean square displacement (MSD) versus time plots of representative trajectories for C-terminally Halo tagged YhdP. (C) Representative brightfield and fluorescent images are shown for C-terminally Halo tagged YhdPwith *dedD* removed (Δ*dedD*). (D) MSD versus time plots of representative trajectories for C-terminally Halo tagged YhdP with *dedD* removed (Δ*dedD*). (E) Representative brightfield and fluorescent images are shown for C-terminally Halo tagged DedD (C) strains. (F) Fluorescence profiles, depicted as demographs, of C-terminally Halo tagged YhdP in a Δ*dedD* background. Data are split between dividing (upper) and non-dividing (lower) cells. Averaged cell shapes for corresponding demographs are also shown.

**Supplementary Figure 6.**
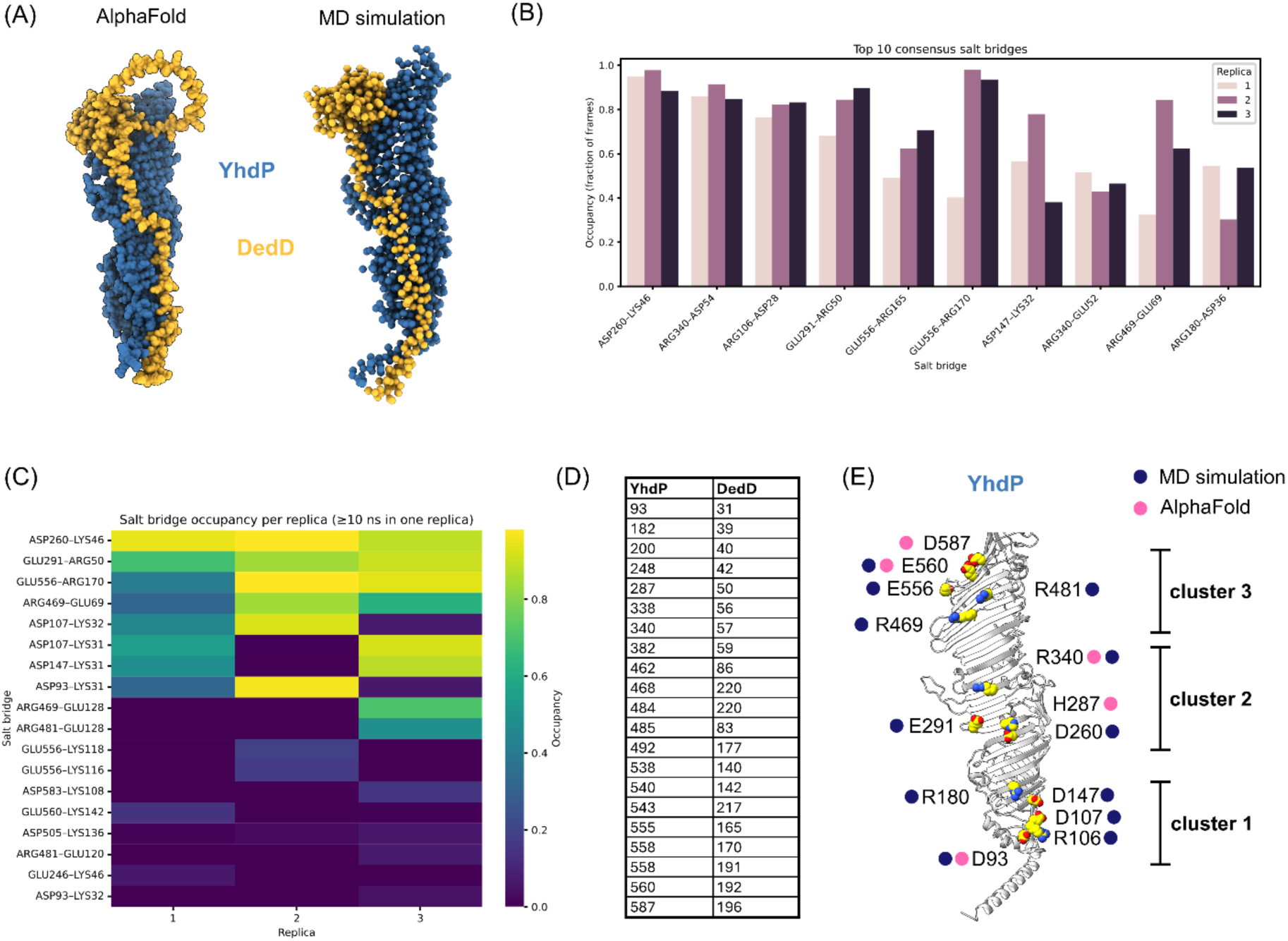
Molecular Dynamics simulations and AlphaFold predicted key salt bridge forming residues for the YhdP-DedD complex. (A) Models of the YhdP (residue 1-605) and DedD complex from AlphaFold and coarse grained MD simulation. In the AlphaFold prediction, the C-terminal region of the DedD IDR extends away from the rest of the protein, whereas during MD simulations it collapses back onto itself without forming contacts with YhdP. (B,C) Salt bridges formed during the MD simulations have been analysed and ranked based on their occupancy, by fraction of frames (B) or salt bridge occupancy duration (C). (D) A list of residues on YhdP and DedD that are predicted to form hydrogen bonds in the AlphaFold model. (E) Schematic diagram of YhdP shows the residues that are investigated by mutagenesis experiments in this study. Residues are coloured based on whether they form salt bridges during MD simulation (blue) and/or in the AlphaFold model (pink). These residues are clustered into three based on their relative locations on YhdP.

**Supplementary Figure 7.**
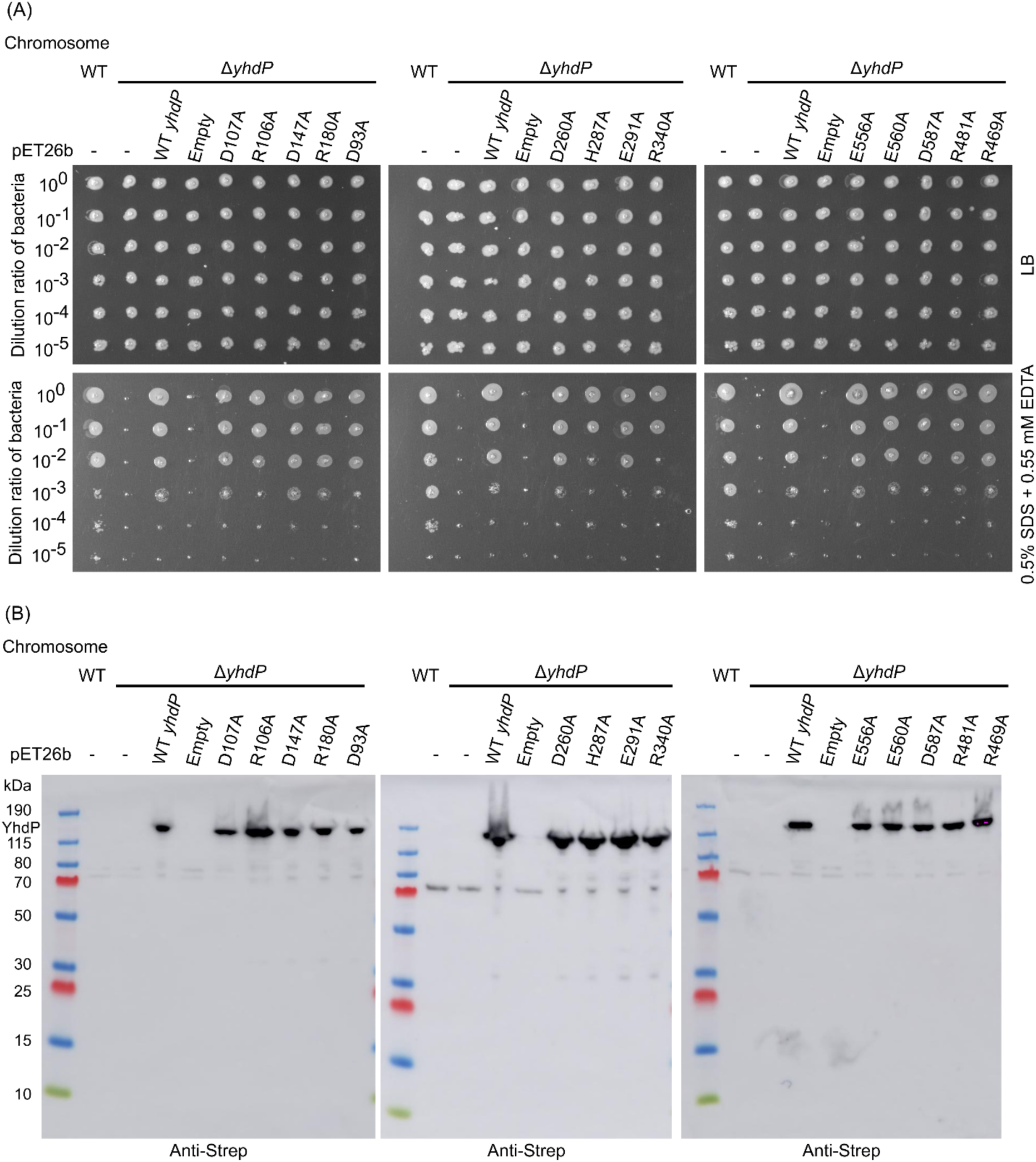
All YhdP single mutants express and function the same as wild-type YhdP. (A) Complementation assays of YhdP single mutants. 10-fold dilution series of indicated strains were spotted on LB agar plates with or without SDS+EDTA. (B) Expressions of these mutants were examined on anti-Strep western blot. WT *yhdP*: Wild-type YhdP with StrepII tag; Empty: empty pET26b vector. Representative images are shown.

**Supplementary Figure 8.**
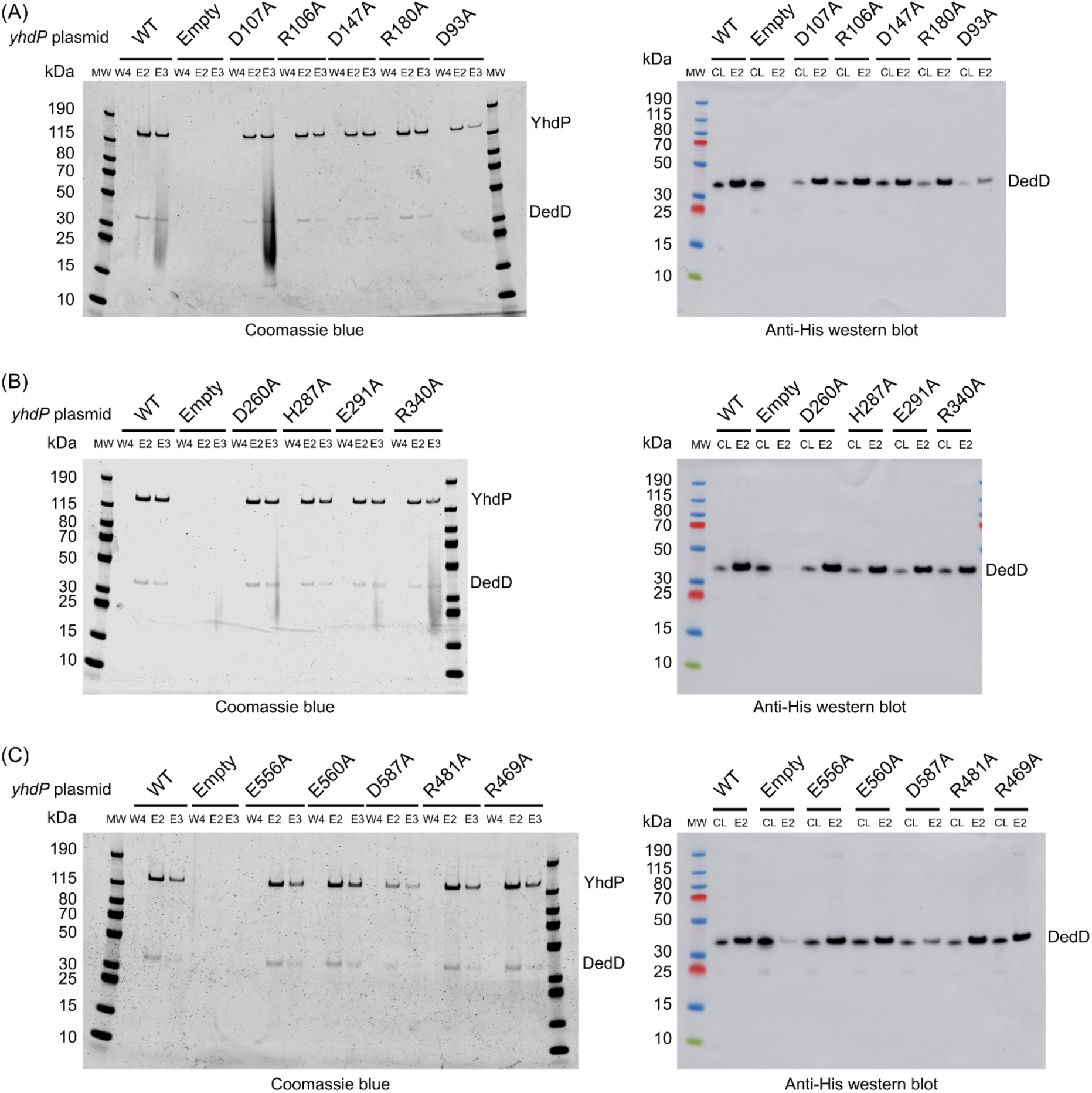
Pull-down of DedD via YhdP single mutants. WT and StrepII-tagged single YhdP mutants were co-expressed with the His-tagged full-length DedD construct. Mutants from cluster 1, 2 and 3 (see Fig. S6A) are shown in panel (A), (B) and (C), respectively. Samples from affinity chromatography were analysed by SDS-PAGE followed by Coomassie-blue staining and anti-His western blot. CL: Whole cell lysate; W4: Final wash; E2: Second elution; E3: Third elution. WT: Wild-type YhdP with StrepII tag; Empty: empty pET26b vector.

**Supplementary Figure 9.**
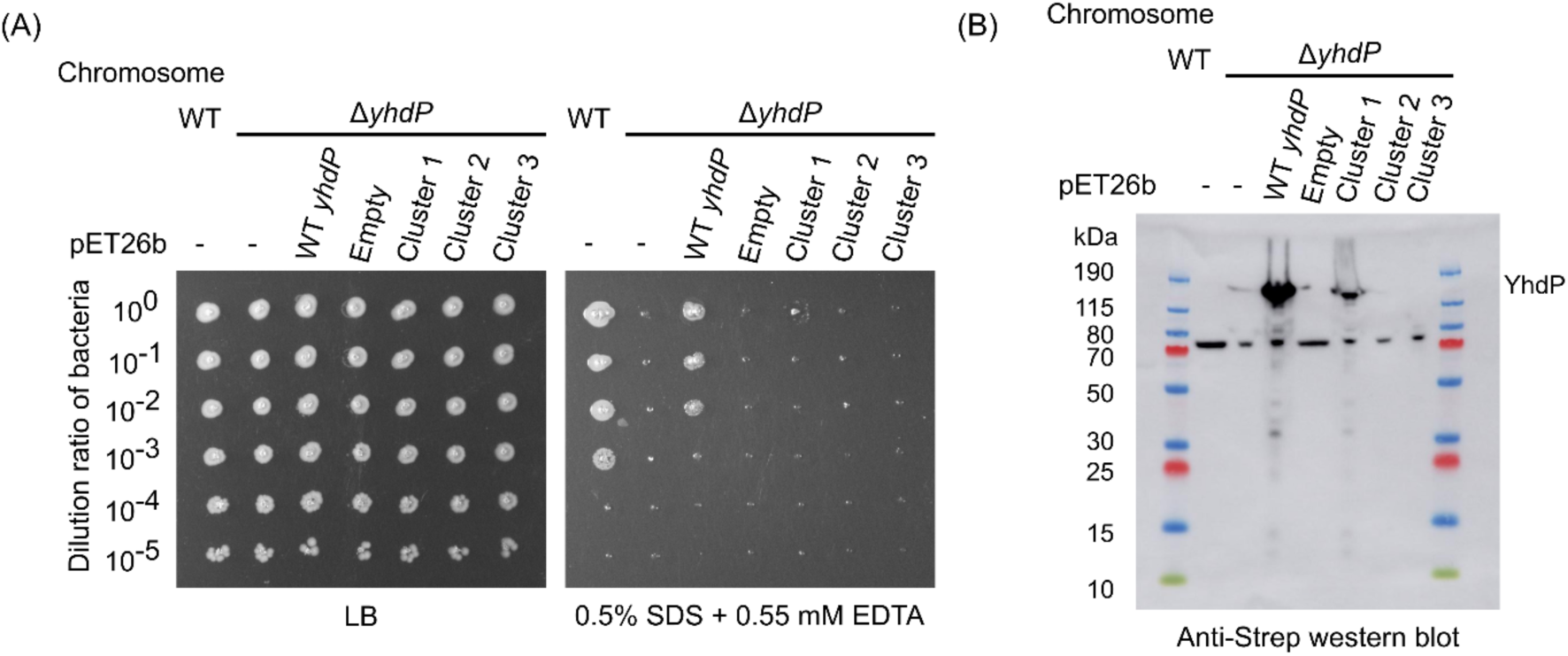
Expression test and pull-down experiments of YhdP cluster mutants. (A) Complementation assays of YhdP cluster mutants. 10-fold dilution series of indicated strains were spotted on LB agar plates with or without SDS+EDTA. Representative images are shown. (B) Expressions of these mutants were also examined on anti-Strep western blot.

**Supplementary Figure 10.**
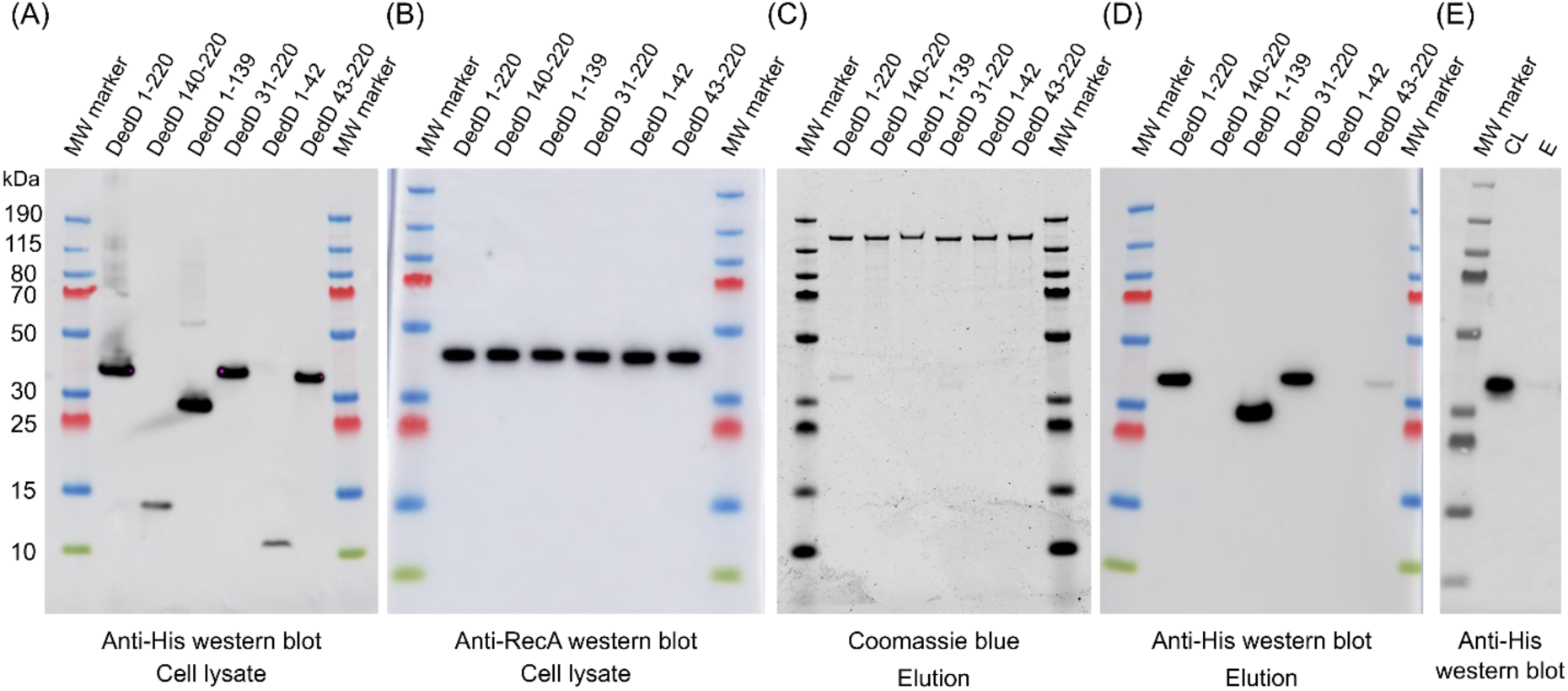
Pull-down of DedD mutants via YhdP. (A) Anti-his western blot against whole cell lysates to detect expression level of DedD mutants. (B) Anti-RecA western blot as a loading control. (C) Uncropped Coomassie blue stain gels of WT YhdP co-expressed alongside each of the DedD constructs. (D) Anti-his western blot against eluted DedD WT and mutants pulled down via YhdP. (E) Control western blot to confirm that His-tagged full-length DedD does not non-specifically bind and elute from Strep resin. CL: cell-lysate; E: second elution.

**Supplementary Figure 11.**
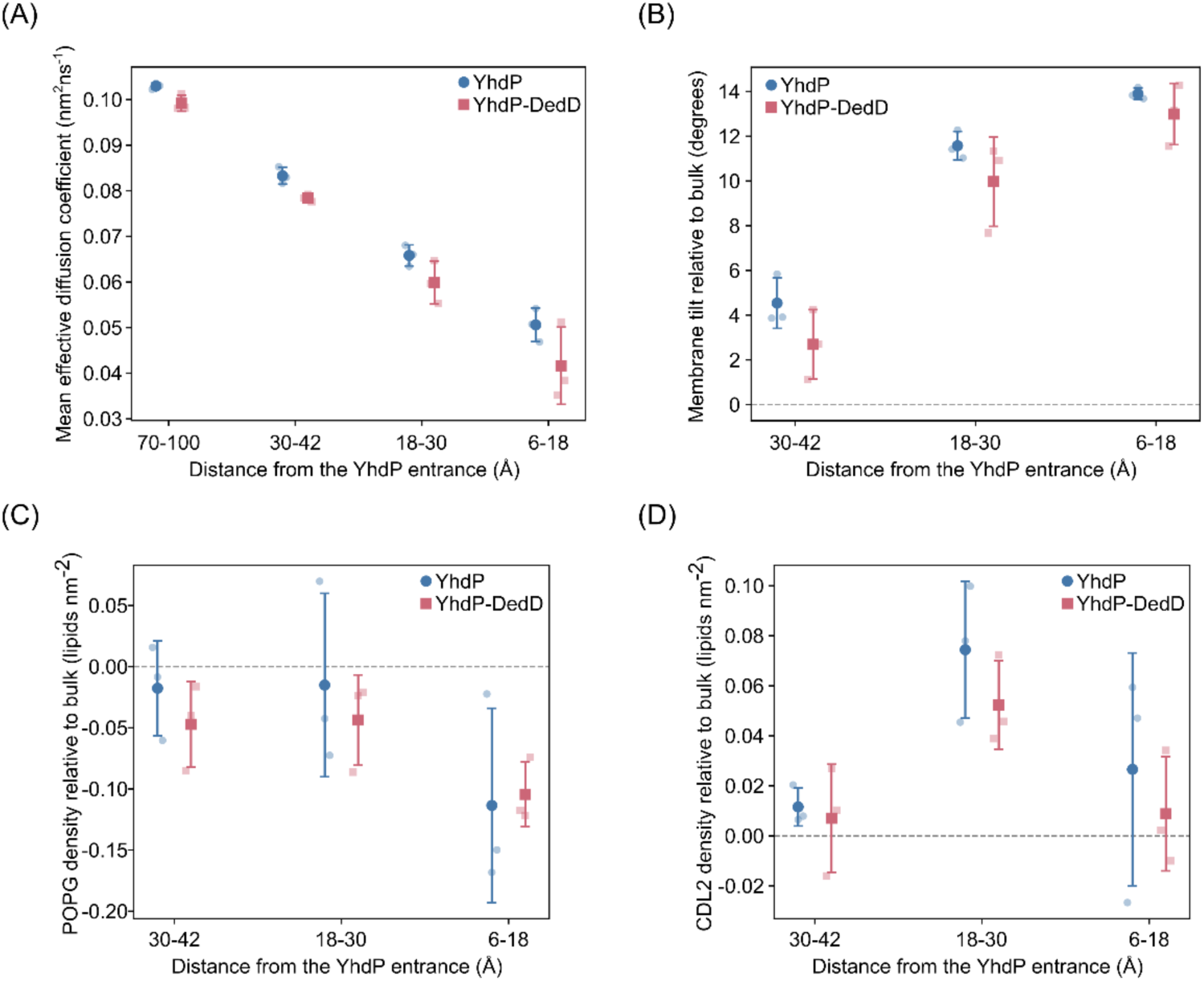
Molecular dynamics simulations study the inner membrane environment of YhdP entry in the absence or presence of DedD. (A) At a 5 ns lag, effective lipid mobility was measured from bulk to 30-42 Å, 18-30 Å and finally 6-18 Å. Mean effective diffusion decreased from 0.103 nm² ns⁻¹ in the bulk membrane to 0.051 nm² ns⁻¹ within 6–18 Å of the entry in the YhdP-only simulations, and from 0.099 to 0.042 nm² ns⁻¹ in the presence of DedD. (B) Normal tilt relative to bulk increases strongly as the entry is approached. The effect is smallest at 30-42 Å, larger at 18-30 Å and the largest at 6-18 Å. (C,D) POPG and CDL2 density is shown relative to each trajectory’s own bulk membrane.

